# Role of GABA and NMDA receptors in shaping cortical timescales and large-scale network dynamics

**DOI:** 10.64898/2026.05.11.723797

**Authors:** Ana Antonia Dias Maile, Oliver Kohl, Eduard Ort, Monja I Froböse, Hannah Kurtenbach, Markus Butz, Elisabeth Schreivogel, Alfons Schnitzler, Esther Florin, Gerhard Jocham

## Abstract

Cortical brain regions integrate information across different timescales, ranging from fast sensory processing to longer integration windows, allowing cognitive functions like working memory. At the large-scale, brain regions organize into transient network states that rapidly switch over time and similarly contribute to cognition. Both cortical timescales and large-scale network dynamics are proposed to be determined by the balance between recurrent synaptic excitation and GABAergic inhibition. Here, we pharmacologically manipulated synaptic transmission at GABA_A_ and NMDA receptors in 60 healthy male participants and acquired resting-state magnetoencephalography. Neuronal timescales followed a hierarchical gradient with shorter timescales in early sensory regions. Increasing GABAergic activity prolonged neuronal timescales across cortical regions. This effect was most prominent in the frontal default mode and in the dorsal attention network. Notably, dynamic network analyses revealed that the occurrence probability of the frontal default mode network increased, whereas the occurrence of the dorsal attention network was reduced. NMDA receptor modulation resulted in no significant changes. Together, these findings provide causal evidence that GABAergic inhibition is a key regulator of cortical temporal organization, linking microscale synaptic mechanisms to neuronal timescales and network dynamics that support diverse cognitive function.

## Introduction

Neurons throughout the brain process information over multiple timescales (Murray et al., 2014). This enables complex perception and cognition, such as following the storyline of a movie: In primary sensory regions, short timescales allow the fast detection of perceptual changes from frame to frame, whereas longer timescales in association regions allow integration of this information into coherent scenes and narratives (Baldassano et al., 2017; Çatal et al., 2023; Honey et al., 2012; Lerner et al., 2011). The spatial distribution of neuronal timescales follows a hierarchical gradient across the cortex, with short timescales in early sensory areas and increasingly longer timescales in higher-order association areas (Murray et al., 2014; Shafiei et al., 2023; Wang, 2020), a pattern that has been replicated across different species and tasks (Cusinato et al., 2023; Halgren et al., 2021). This gradient and its dynamic adaptation to cognitive demands play a fundamental role in information processing and working memory (Bernacchia et al. 2011; Gao et al., 2020; Ito et al., 2020; Lendner et al. 2024; Manea et al., 2024). In addition to these local dynamics, distant brain regions also spontaneously organize into short-lived, transient network states with distinct spectral signatures (Vidaurre et al. 2018) that likewise support cognitive processes on different timescales (Quinn et al., 2018; Gohil et al., 2024; Rossi et al., 2023; Rossi et al., 2024). This ability of the brain to explore a diverse repertoire of large-scale network configurations can be seen as sequential activation of recurring, metastable states (Rabinovich et al., 2008; Tognoli & Kelso, 2014).

Neuronal timescales and metastable resting-state dynamics are proposed to emerge from the balance between glutamatergic excitation and GABAergic inhibition (Excitation-inhibition-balance, or short, E/I balance; Abeysuriya et al., 2018; Gao et al., 2017; Murray et al., 2014; Pascoa Dos Santos & Verschure, 2025; Wang, 2020). Areas at the top of the cortical hierarchy, like the prefrontal cortex are rich in recurrent connections endowed with NMDA glutamate receptors (Burt et al., 2018). Owing to their long time constant, they enable persistent neural activity and information integration over long timescales critical for working memory and decision making (Wang et al., 2008; Wang, 1999, 2002, 2008). In contrast, NMDA receptor expression is less pronounced in early sensory areas (Burt et al., 2018). Consequently, neural activity there fluctuates on shorter timescales and closely tracks the current sensory input (Baldassano et al., 2017; Honey et al., 2012). Recurrent excitation is balanced by shorter GABAergic feedback inhibition, which displays a similar hierarchical gradient (Burt et al., 2018). Despite these findings, it is not known how NMDA-and GABA-mediated activity causally shapes neuronal timescales across cortical regions. It further remains unclear whether different large-scale cortical networks have distinct neuronal timescales and how transitions between network states are governed by changes in transmission at GABA and NMDA receptors.

In the present study, we examine how NMDA and GABA_A_ receptors influence both neuronal timescales across cortical regions and the dynamical organization of large-scale cortical networks. To this end, we increased activity at NMDA and GABA_A_ receptors by administering the partial NMDA receptor agonist D-cycloserine and the benzodiazepine receptor agonist lorazepam, respectively, and examined their effects on the hierarchical organization of cortical timescales and on large-scale network dynamics during rest. Given the slower kinetics of NMDA and GABA receptors relative to the predominant AMPA-mediated excitation (Angulo et al., 1999; Gupta et al., 2000; Sah et al., 1990), we hypothesized that both manipulations would prolong neuronal timescales. In brief, we find that increased GABAergic activity prolongs neuronal timescales across the cortex, differentially modulates timescales across dynamic large-scale cortical networks, and alters network dynamics by biasing transitions toward the frontal default mode network. Modulation of NMDA receptor activity did not significantly alter neuronal timescales or large-scale network dynamics.

## Results

We examined how NMDA and GABA_A_ receptors influence neuronal timescales across cortical regions and the dynamical organization of large-scale cortical networks by analyzing five minutes of open-eyed resting-state data. To this end, we collected magnetoencephalography (MEG) recordings from 60 healthy participants under the influence of lorazepam (1 mg), D-cycloserine (250 mg), or placebo in a double-blind, within-subject, crossover design. Lorazepam is a benzodiazepine receptor agonist that binds allosterically at GABA_A_ receptors (Riss et al., 2008; Tallman et al., 1978). D-cycloserine is a glutamatergic partial NMDA agonist acting at the glycine-binding site (Thomas et al., 1988). Three participants were excluded due to excessive noise in the MEG data, resulting in a final sample of 57 participants.

### Timescales follow a hierarchical cortical gradient

Neuronal timescales reflect the decay rate of the autocorrelation function of a neural signal (i.e., the signal’s decreasing self-similarity across time). Timescales were estimated per cortical region of the Desikan-Killiany atlas (34 parcels per hemisphere; Desikan et al., 2006) based on the aperiodic knee frequency fitted with the FOOOF toolbox (version 1.0.0; Donoghue et al., 2020; Gao et al., 2020, see Methods for more details). As an initial validation, we first assessed the spatial similarity between the estimated timescales under placebo and the publicly available intrinsic timescale map (Glasser et al., 2016; Shafiei et al., 2023). We observe a strong correlation of *r_s_* = 0.91, supporting the validity of our estimates (*p* < 0.001; permutation test preserving the spatial autocorrelation; see Methods).

As it has been indicated that timescales follow a hierarchical gradient (Gao et al., 2020; Murray et al., 2014; Shafiei et al., 2023; Wang, 2020), we tested how timescales relate to the T1w/T2w ratio, an MRI-based index for myelinization commonly used as a proxy for a cortical region’s position on the cortical hierarchy, with lower ratios indicating higher hierarchical position (Glasser et al., 2016). In agreement with this, timescales under placebo - but also under both drug conditions - were strongly negatively correlated with the T1w/T2w ratio (all *r_s_* <-0.79, *p* = 0.002; permutation test preserving the spatial autocorrelation; see Methods and Figure 1). Thus, we confirm that neuronal timescales increase with higher position in the cortical hierarchy. Importantly, this gradient was preserved under pharmacological increase of GABA_A_ and NMDA receptor-mediated transmission.

**Fig. 1:**
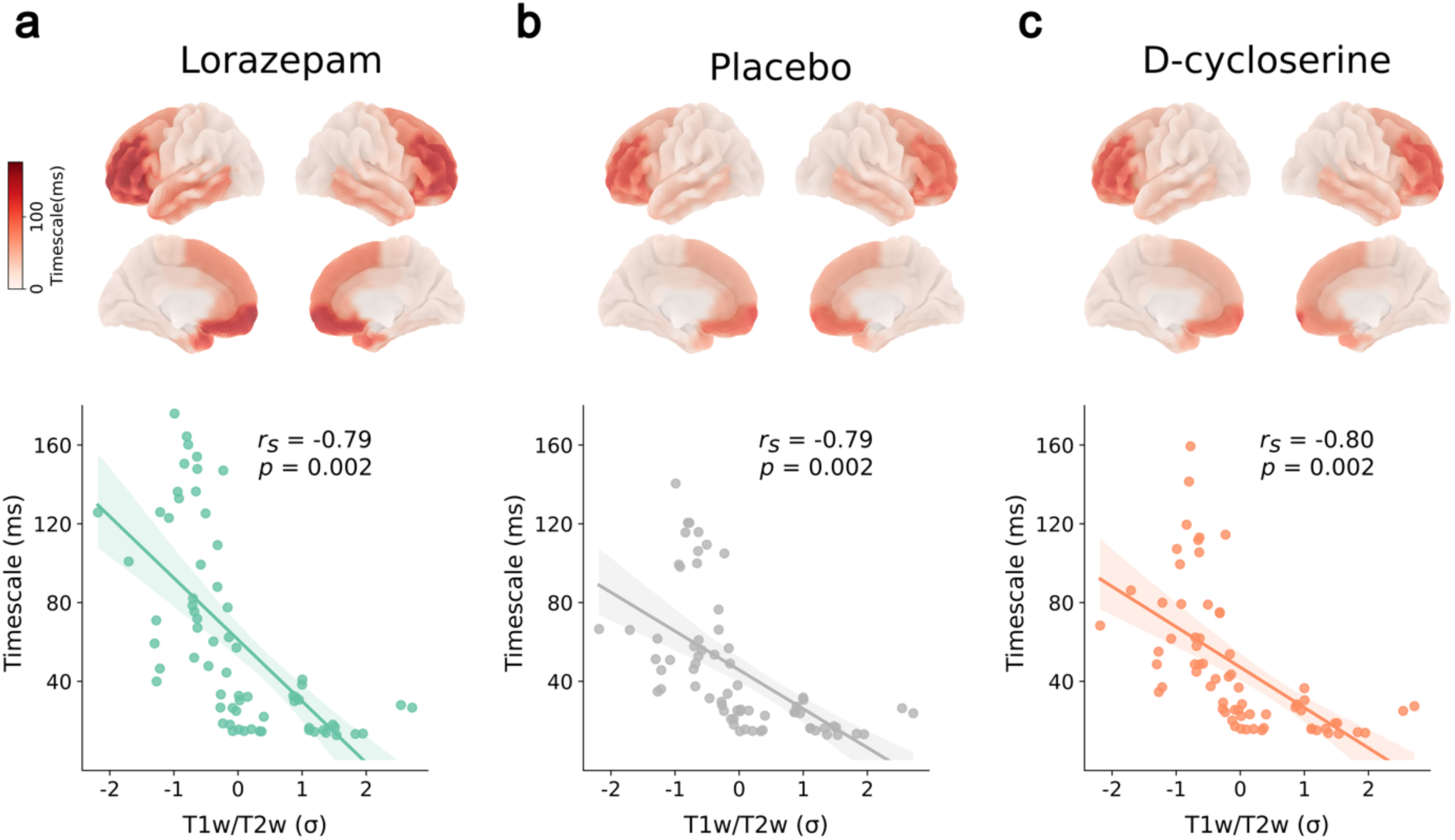
Hierarchical gradient of neuronal timescales. Timescales for lorazepam **(a)**, placebo **(b)**, and D-cycloserine **(c). Top row:** Distribution of neuronal timescales in milliseconds (averaged across participants) across the cortex. **Bottom row:** Timescales of all 68 cortical areas (averaged across participants) as a function of T1w/T2w ratio (z-scored), an index of the hierarchical position of a cortical area. Shades around the regression line represent the 95% confidence interval. *rS* = Spearman correlation coefficient. Spatial autocorrelation was preserved for the computation of *p*-values.

### Pharmacological modulation of neuronal timescales

Recurrent excitation at NMDA receptors and GABAergic feedback inhibition is thought to shape temporal integration of neurons (Wang, 2020). In line with this, lorazepam, compared with placebo, significantly increased neuronal timescales across 48 of 68 parcels in both hemispheres (Figure 2a and Supplement 1; see Supplement 2 for drug effects on all aperiodic parameters; two-tailed paired-sample permutation t-test, all *t* > 2.14, *p* < 0.044 with FDR correction for multiple comparisons). In contrast, D-cycloserine resulted in no significant changes in neuronal timescales (Figure 2b).

**Fig. 2:**
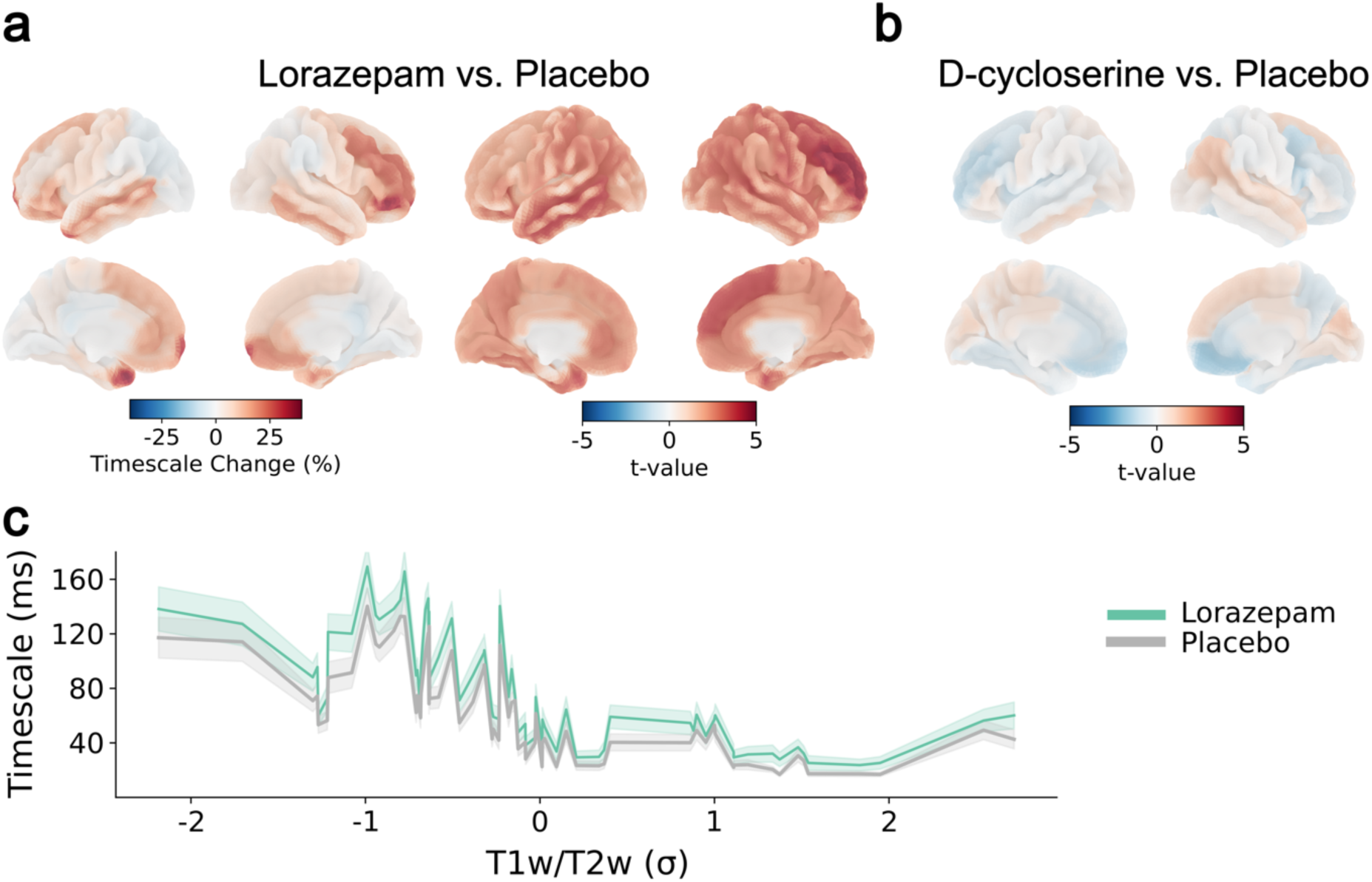
Drug effects on neuronal timescales and their hierarchical gradient. **a** Left panel: percent change of timescales under lorazepam compared to placebo. Right panel: *t*-statistic on neuronal timescales, permutation test of the effect of lorazepam compared to placebo, projected onto the cortical surface. **b** *t*-statistic on neuronal timescales, permutation test of the effect of D-cycloserine on neuronal timescales compared to placebo, projected onto the cortical surface. D-cycloserine did not significantly modulate neuronal timescales. **c** Cortical hierarchy (operationalized via the z-scored T1w/T2w ratio) of neuronal timescales as a function of drug condition (lorazepam or placebo). Shades depict the 95%-confidence intervals.

Previous work suggests a relation between a brain region’s level of GABA receptor expression and its neuronal timescale, above and beyond the regions’ hierarchical position (Gao et al., 2020). Therefore, we next examined whether the effects of lorazepam were independent of the position of the cortical parcels in the cortical hierarchy. A hierarchy-dependent effect of lorazepam would be evident from a change in the slope of the regression line between cortical hierarchy and neural timescales (Figure 1a). To this end, we implemented a linear mixed-effects regression on neuronal timescales, including the predictors cortical hierarchy (T1w/T2w ratio), drug condition, and their interaction. In line with the previous analyses, the T1w/T2w ratio had a significant main effect on neuronal timescales (*β* =-3.06, *SE* = 0.17, *t*_75880_ =-17.64, *p* < 0.001; supplement 3), indicating that increased myelination was associated with shorter timescales. Moreover, lorazepam showed a significant positive main effect (*β* = 0.39, *SE* = 0.15, *t*_5146000_ = 2.55, *p* = 0.011), confirming that lorazepam prolongs neuronal timescales. There was also a trend for an interaction between drug condition and T1w/T2w ratio (*β* =-0.19, *SE* = 0.11, *t*_11190000_ =-1.75, *p* = 0.081; Figure 2c). This trend points towards an increased hierarchical gradient under lorazepam, likely due to a larger increase of neuronal timescales in frontal cortex compared to occipital cortex (Figure 2a). Importantly, the increase in timescales under lorazepam, but also the trend-wise interaction of lorazepam with the cortical hierarchy, are independent of the significant effects of lorazepam on various control measures we collected (i.e., alertness, calmness, contentedness, anxiety, visuomotor abilities, and vital parameters; see Supplement 4 and 5). Consistent with the results obtained from the permutation t-test across cortical regions (Fig. 2b), D-cycloserine had neither a significant main effect (*β* =-0.08, *SE* = 0.15, *t*_9952000_ =-0.54, *p* = 0.586) nor an interaction with the cortical hierarchy (*β* = 0.08, *SE* = 0.11, *t*_11180000_ = 0.71, *p* = 0.477), nor on control parameters (see Supplement 4).

In sum, neuronal timescales increase along the cortical hierarchy and are further prolonged by lorazepam. While this effect appears largely independent of the hierarchical gradient, there is a trend toward stronger modulation in the frontal cortex, leading to a steeper cortical gradient.

### Dynamic cortical large-scale networks exhibit varying timescales

Having established that local neuronal timescales vary systematically across the cortical hierarchy and are increased by lorazepam, we next examined timescales of dynamic large-scale cortical networks. To this end, we applied a Time-Delay Embedded Hidden Markov Model (TDE-HMM) to the parcellated MEG time series to extract spectrally and temporally resolved large-scale cortical networks. The resulting eight networks (Figure 3) closely resemble previously reported TDE-HMM-derived cortical networks (Gohil et al., 2024; Kohl et al., 2025; Rossi et al., 2023; Rossi et al., 2024; Vidaurre et al., 2018). All states were expressed in every participant, with fractional occupancies ranging from 1 % to 48 %. Fractional occupancy refers to the percent of the entire recording time the brain spent in the respective state. This demonstrates that no single state dominated individual recordings and that each network contributed to ongoing cortical dynamics.

**Fig. 3:**
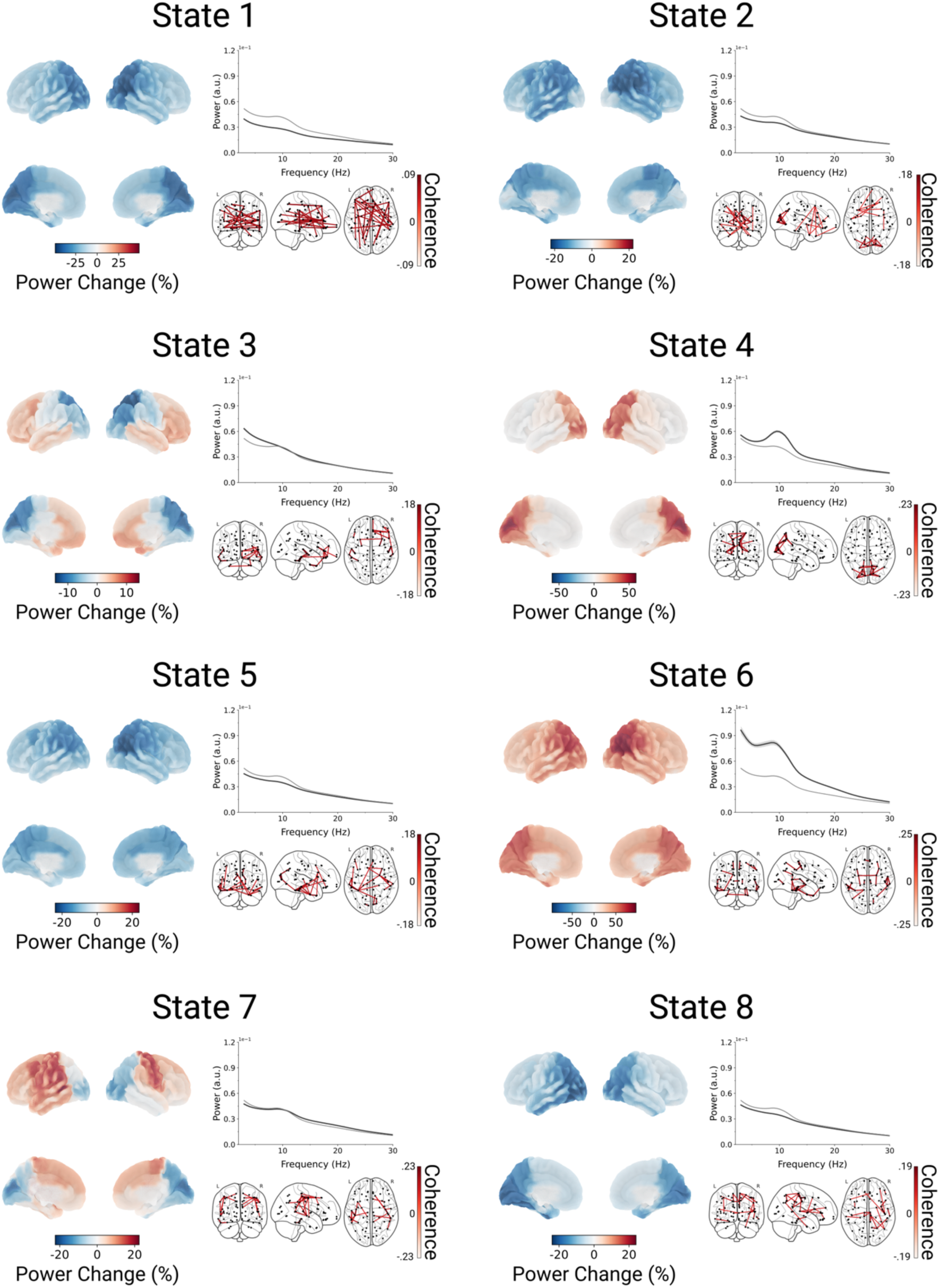
Spectral characterization of dynamic large-scale cortical networks. For each state, three representations are presented, averaged across all participants in the placebo condition. **Left:** Cortical maps of state-specific 3-30 Hz power changes relative to the time-averaged power across states. **Top right:** Whole-brain average power spectrum for each state (black) compared with the whole-brain average spectrum across all states (grey). **Bottom right:** State-specific coherence networks in the 2–30 Hz range, thresholded at the 98th percentile to highlight the strongest functional connections.

Next, we estimated state-specific timescales from each state’s power spectrum, using the same procedure as reported above for time-averaged spectra. These state-specific timescales were then compared to each participant’s global timescales using General Linear Models (GLMs) to determine whether neuronal timescales systematically varied with the occurrence of specific cortical networks. Visits to two states were associated with significant increases in timescales compared to the global timescales across multiple cortical regions (State 1 and State 3: all *t*_55_ > 4.79, *p* < 0.001; max change = +5.46% to +21.7%; Figure 4a). In contrast, for the remaining states, the time-scales predominantly decreased significantly (State 2, 4, 5, 7, and 8: all *t*_55_ < - 2.18, *p* < 0.030; max change =-2.22% to –27.87 %) or the pattern was mixed (State 6: six parcels increased: all *t*_55_ > 3.11, *p* < 0.001; 3 parcels decreased: all *t*_55_ <-2.75, *p* < 0.006; max change =-15.10 % to +8.99 %, Figure 4a). This suggests that large-scale cortical networks operate on distinct timescales, indicating a flexible temporal organization of cortical processing.

**Fig. 4:**
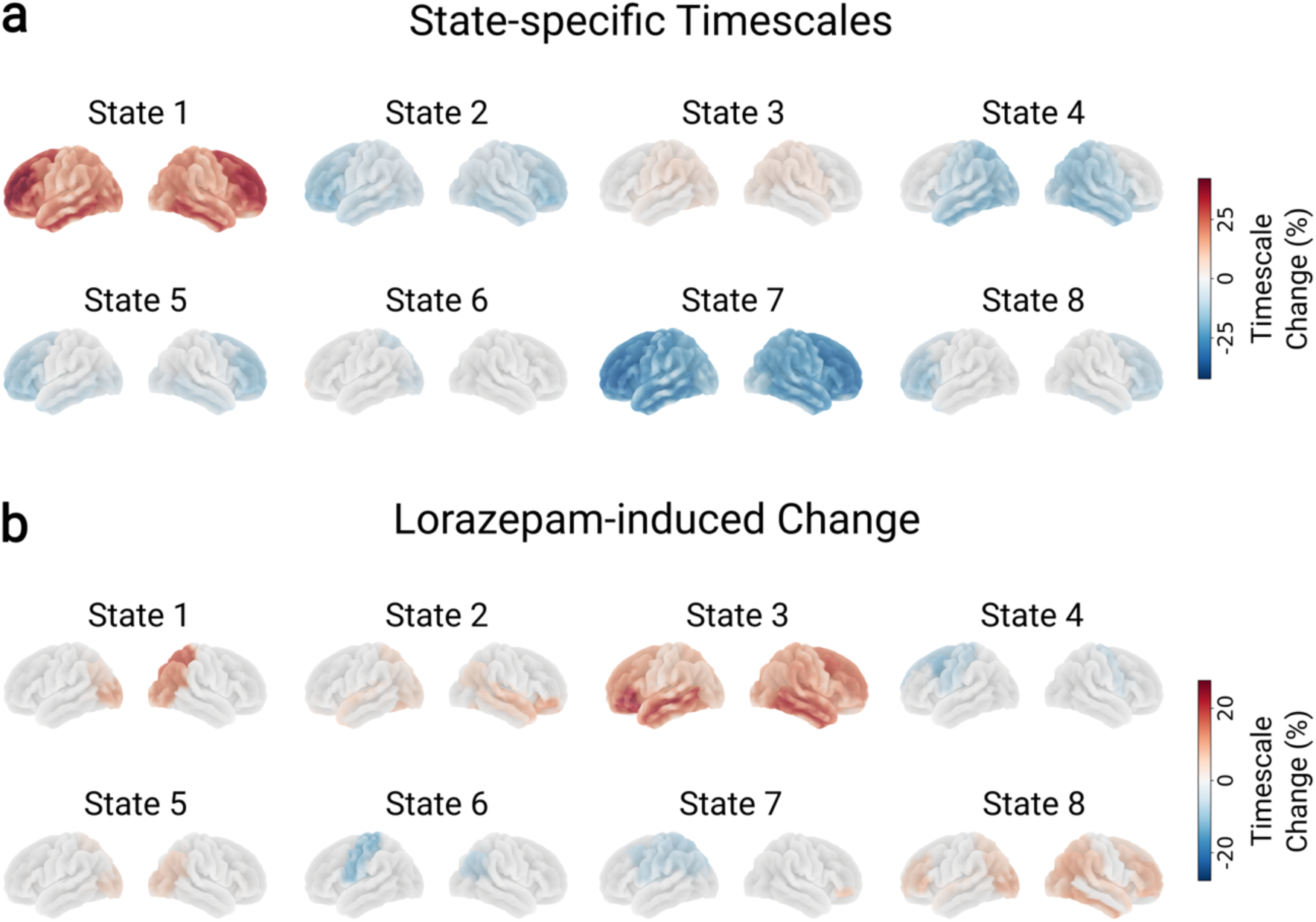
Timescales vary across cortical networks and are differentially modulated by lorazepam. **a** Percent change in neuronal timescales during network occurrences relative to the time-averaged timescale across states. **b** Percent change in neuronal timescales under lorazepam relative to placebo. Only parcels with significant effects are shown (*p* < 0.05, FDR-corrected). Significance was assessed for each parcel and state using within-subject GLMs with permutation testing, comparing state-specific timescales with the time-averaged timescale (**a**) or between lorazepam and placebo (**b**). p-values were FDR-corrected across parcels and states.

### Pharmacological modulation of network-specific time scales

The above analyses demonstrate that large-scale cortical networks operate on different timescales. Next, we investigated how our pharmacological intervention modulates these state-specific timescales. As D-cycloserine showed no significant effects on neuronal timescales, the subsequent analyses focus exclusively on lorazepam. GLMs comparing state-specific timescales between the lorazepam and placebo condition revealed significantly increased timescales during visits to State 1 (*n* = 12/68 significant parcels; max increase = 17.52%, *t*_55_ = 4.75, *p* < 0.001), State 2 (*n* = 21/68 significant parcels; max increase = 6.38%, *t*_55_ = 3.9, *p* < 0.001), State 3 (*n* = 59/68 significant parcels; max increase = 24.02, *t*_55_ = 5.34, *p* < 0.001), State 5 (*n* = 10/68 significant parcels; max increase = 6.39, *t*_55_ = 4.15, *p* < 0.001), and State 8 (*n* = 34/68 significant parcels; max increase = 9.8, *t*_55_ = 5.28, *p* < 0.001; Figure 4b), whereas for the remaining states timescales significantly decreased (all *t*_55_<-1.86, *p* < 0.020; max decrease =-5.47% to-13.82%). This pattern indicates that lorazepam exerts network-specific effects on neuronal timescales, such that its influence on a given cortical region varies depending on which large-scale network is currently active.

The lorazepam-related increase in neuronal timescales across most networks is consistent with the widespread prolongation of cortical timescales observed in our time-averaged region-wise analysis. Among the networks, the strongest lorazepam-induced increases of neuronal timescales were observed for State 3 (∼24%) and State 1 (∼17.5%), which have been previously linked to the frontal default mode network (DMN; Gohil et al., 2024; Rossi et al., 2023; Rossi et al., 2024; van Es et al., 2025; Vidaurre et al., 2018) and the dorsal attention network (DAN; Baker et al., 2014; van Es et al., 2025), respectively. Despite these strong increases, repeated measures correlations across all participants and parcels indicated small but significant associations (all *r* < 0.22) between global and state-specific timescales for all states except the DAN (State 1; see Supplement 6). In contrast, frontal DMN (State 3) timescales exhibited the strongest coupling with global timescales (*r* = 22, *p* <.0.001). Together, these findings indicate that the frontal DMN is an important driver of the global prolongation of neuronal timescales across the cortex under lorazepam.

### Lorazepam shifts cortical dynamics toward the frontal default mode network

After identifying that neuronal timescales vary across cortical networks and are differentially modulated by lorazepam, we next examined whether lorazepam also alters large-scale network dynamics. Network dynamics were characterized using fractional occupancy (the overall proportion of time a network is active) and complementary temporal metrics: state lifetime (the average duration of continuous visits); interval time (the time between successive visits); and state rate (the frequency of network entry per second). GLMs comparing participants’ fractional occupancies between the placebo and lorazepam conditions revealed a significant increase for the frontal DMN (State 3; *t*_55_ = 4.33, *p* < 0.001) and a significant decrease for the DAN (State 1; *t*_55_ =-3.08, *p* = 0.020; Figure 5a) under lorazepam. Analyses of lifetimes, interval times, and state rates indicated that the enhanced fractional occupancy of the frontal DMN (State 3) was driven by increases in both state lifetime (*t*_55_ = 2.84, *p* = 0.040; Figure 5b) and state rate (*t*_55_ = 3.97, *p* < 0.001, Figure 5d), whereas the reduction in DAN (State1) occupancy was underpinned by shorter state lifetimes (*t*_55_ =-3.09, *p* = 0.020, Figure 5b). Interval times were not modulated by lorazepam (Figure 5c).

**Fig. 5:**
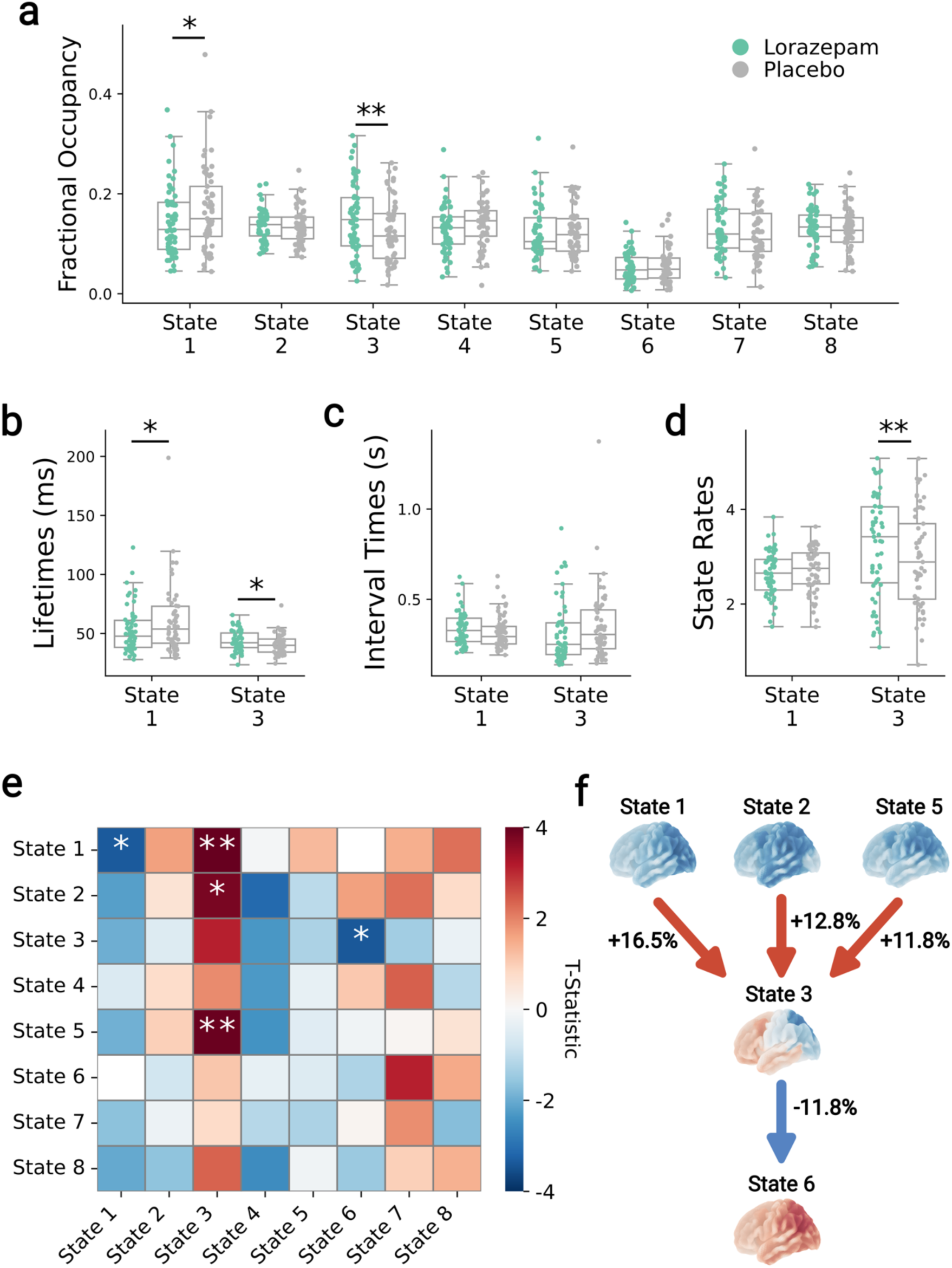
Lorazepam modulates large-scale cortical network dynamics. **a** Lorazepam–placebo contrasts of fractional occupancy across all states. Each dot represents an individual participant. Lorazepam reduced the fractional occupancy of State 1 (dorsal attention network, DAN) and increased that of State 3 (frontal default mode network, DMN). **b-d** Lorazepam-placebo contrasts of state lifetimes (**b**), interval times (**c**), and state rates (**d**) for the DAN and frontal DMN. Dots indicate individual participants. **e** Lorazepam-placebo contrasts of state transition probabilities. Heatmap colours represent *t*-statistics from within-participant GLMs. Rows denote the current state and columns the subsequent state. Lorazepam reduced DAN self-transitions, increased transitions from the DAN and other networks into the frontal DMN, and reduced transitions out of the frontal DMN. **f** Schematic summary of significant lorazepam-induced transition changes involving the frontal DMN. Values indicate percentage change relative to placebo, with arrows denoting transition direction. Statistical significance in **a-e** was assessed using maximum *t*-statistic permutation tests, correcting for multiple comparisons across states or transitions. Asterisks indicate significance: *p* < 0.050 = *, *p* < 0.010 = **.

To further detail the mechanisms underlying the lorazepam-related increase in frontal DMN occupancy, we compared state-transition probabilities between the lorazepam and placebo conditions. This analysis tests whether the temporal progression from one network state to another differs as a function of drug condition. GLMs combined with maximum *t*-statistic permutation tests revealed that lorazepam significantly reduced DAN self-transitions (State 1; *t*_55_ =-3.4, *p* = 0.050, Figure 5e), consistent with the observed reductions in occurrence probabilities and lifetimes. Furthermore, under lorazepam, the DAN (State 1; *t*_55_ = 4.05, *p* = 0.007), State 2 (*t*_55_ = 3.83, *p* = 0.010), and State 5 (*t*_55_ = 4.01, *p* < 0.007) were significantly more likely to transition into the frontal DMN (State 3), while transitions out of the frontal DMN were significantly reduced (State 6: *t*_55_ = −3.45, *p* < 0.040; Figure 5e,f). These findings suggest that lorazepam increases transitions into and decreases transitions out of the frontal DMN, thereby increasing the time the brain spends in that state.

### Robustness Analyses

TDE-HMM analyses require a priori specification of state number, as standard model selection metrics do not identify a clear optimum (Gohil et al., 2026). We therefore repeated all key analyses using models with 10 and 12 states. Results were similar across different numbers of states for state-specific timescales, lorazepam effects on timescales (Supplement 7), and lorazepam-effects on network fractional occupancies (Supplement 8). Similarly, re-running the TDE-HMM using a different preprocessing pipeline (Supplement 9) also produced comparable states, indicating that the main findings are not dependent on state number selection or specific preprocessing choices.

## Discussion

The present study examined the causal role of GABAergic and NMDA-mediated neurotransmission in shaping neuronal timescales and large-scale network dynamics. Our results demonstrate that enhancing GABAergic inhibition via lorazepam produced a widespread prolongation of neuronal timescales across the cortex, with a trend toward stronger effects in frontal regions, indicative of a steepened hierarchical gradient. These timescale changes were most pronounced during activation of the frontal DMN and DAN, and were accompanied by opposing shifts in network recruitment, with lorazepam increasing the probability of frontal DMN occurrence while reducing that of the DAN. In contrast, facilitating NMDA receptor transmission via D-cycloserine had no significant effect on cortical timescales. Together, these findings provide causal evidence that microscale synaptic inhibition directly shapes both local neuronal timescale organization and macroscale network dynamics, thereby linking synaptic activity to the hierarchical structure of cortical processing.

In the present study, we replicate observations that neuronal timescales follow a cortical hierarchy, increasing from primary sensory to association areas (Gao et al., 2020; Murray et al., 2014). We further extended these findings by demonstrating that this hierarchy is preserved under elevated transmission at GABA_A_ and NMDA receptors. Higher-order cognitive processes, such as working memory (Gao et al., 2020; Wang et al., 2008; Wang et al., 2013; Wang, 1999) and decision making (Wang, 2002, 2008), strongly depend on recurrent NMDA-mediated excitation. Increased NMDA-mediated recurrent excitation is thought to prolong neuronal timescales, which fits with the observation that, during behavioral tasks, neuronal timescales are similarly expanded (Bernacchia et al. 2011; Manea et al., 2024), while the general hierarchy is preserved (Manea et al., 2024). We hypothesized that pharmacologically increasing transmission at NMDA receptors with D-cycloserine would likewise increase neuronal timescales. However, we observed no significant effects of D-cycloserine. By contrast, and consistent with our hypothesis for GABAergic modulation, lorazepam prolonged neuronal timescales across almost the entire cortex. Glutamatergic and GABAergic receptors exhibit distinct timescales, with AMPA receptors operating on the shortest timescale (∼2 ms) followed by GABA (∼5-10 ms; Gupta et al., 2000) and NMDA (∼50-100 ms; Angulo et al., 1999; Sah et al., 1990) receptors. At rest, the E/I balance is dominated by GABAergic inhibition and AMPA-mediated excitation, whereas NMDA receptor activity has a reduced contribution due to the voltage-dependent Mg^2+^ block (Alberto & Hirasawa, 2010; Espinosa & Kavalali, 2009; Myme et al., 2003; Vargas-Caballero & Robinson, 2004). Although D-cycloserine binds allosterically at the NMDA receptor and thus increases the probability of channel opening, postsynaptic depolarization is still required for NMDA currents to emerge (Thomas et al., 1988). This may explain the absence of significant effects of D-cycloserine on neuronal timescales at rest. This reasoning also explains the direction of the GABAergic effect: lorazepam prolongs neuronal timescales by strengthening slower GABAergic inhibition and reducing faster AMPA-mediated excitation (Chapman et al., 2022; Isaacson & Scanziani, 2011). Consistent with this interpretation, AMPA antagonists modulate resting state periodic brain activity in a manner similar to what has been observed under GABA agonists (Nutt et al., 2015; Routley et al., 2017). Likewise, propofol anesthesia, which is thought to increase GABAergic inhibition, steepens the aperiodic slope indicative of longer neuronal timescales (Colombo et al., 2019; Gao et al., 2017; Waschke et al., 2021).

In the present study, we observed not only an overall prolongation of timescales but also a trend for greater effects in frontal cortical regions. In line with these findings, neuronal timescales correlate with cortical expression patterns of genes encoding GABA_A_ receptors, both along the cortical hierarchy and beyond it (Gao et al., 2020). In sum, increased GABAergic transmission via lorazepam led to a widespread prolongation of neuronal timescales across the cortex, while modulation of NMDA-mediated receptor activity by D-cycloserine had no significant effects.

In addition to local region-wise differences between brain regions, timescales systematically differed across large-scale cortical networks. These networks are thought to play a central role in the organization of information processing in the brain and have been implicated in a wide range of functions, including sensory processing and higher-order information integration (Bressler, 1995; McIntosh, 2000; Mesulam, 1998). In line with the hierarchical gradient of neuronal timescales, states involved in higher-order processing, such as State 1 (DAN) and State 3 (DMN), were characterized by higher intrinsic timescales, relative to global timescales. In contrast, states reflecting sensorimotor or visual networks (Gohil, Kohl, et al., 2024; Quinn et al., 2018), which are more closely tied to bottom-up processing, exhibited shorter timescales. These findings support the notion that the temporal properties of neuronal dynamics support the functional roles of large-scale cortical networks.

Comparisons of network-specific timescales between placebo and lorazepam revealed prolonged timescales for most networks, consistent with our global analysis. The largest increases in timescale occurred in the occipito-parietal areas of the DAN (State 1) and the prefrontal-temporal areas of the frontal DMN (State 3). These cortical areas overlap with regions of high GABA_A_ receptor density (la Fougere et al., 2011; Norgaard et al., 2021), suggesting that regional receptor distribution modulates lorazepam-induced timescale prolongation. Despite this, both the magnitude and direction of lorazepam’s effects varied across resting-state networks and cortical regions. Notably, state-specific spectral power in sensorimotor regions was either increased or decreased by lorazepam depending on current network activation. This pattern suggests that network-specific variability cannot be explained solely by differences in regional recruitment across networks. Instead, the effects of lorazepam on individual cortical regions were not fixed but depended on the present large-scale network activation, indicating state-dependent GABAergic modulation. A plausible explanation is that distinct networks preferentially recruit neuronal subpopulations with differing sensitivity to lorazepam. Benzodiazepines act on multiple GABA_A_ receptor subtypes, which are heterogeneously distributed (Fritschy & Mohler, 1995; Rudolph & Knoflach, 2011) and differentially shape inhibitory gain (Colombi et al., 2024; Patel et al., 2016) and oscillatory dynamics (Heistek et al., 2013). If different networks engage circuits with distinct receptor profiles, this could account for the observed network-specific effects. In support of this, neuronal timescales of the frontal eye field depend on activation of the frontoparietal network, likely reflecting the recruitment of distinct neuronal subpopulations (Soyuhos et al., 2026). In sum, lorazepam effects vary across large-scale cortical networks, highlighting interactions between microscale GABAergic activity and macroscale network dynamics that may be important to consider when investigating pharmacological effects on brain activity.

Analysis of large-scale network dynamics revealed that increasing GABAergic activity via lorazepam modulated not only the intrinsic timescales of the frontal DMN (State 3) and DAN (State 1), but also their temporal expression. Specifically, lorazepam increased the probability of frontal DMN activation, driven by prolonged state lifetimes and higher occurrence rates, while reducing DAN expression, primarily through shortened lifetimes. Analyses of state transition probabilities further suggested that these effects may be underpinned by attractor strength of the frontal DMN, reflected in greater transition rates from multiple networks, including the DAN, into the frontal DMN, and reduced transitions from the frontal DMN into State 6. The opposing modulation on DMN and DAN activity is consistent with their well-established anticorrelation (Baker et al., 2014; Fox et al., 2005; van Es et al., 2025), commonly interpreted as reflecting a balance between internally and externally oriented modes of processing, or task-relevant versus task-irrelevant thoughts (Fox et al., 2005). Taken together, these results suggest that increased GABAergic activity biases large-scale brain dynamics toward internally oriented processing while attenuating responsiveness to external stimuli, consistent with the known sedative effects of lorazepam.

Critically, the observation that pharmacological manipulation of GABAergic activity altered large-scale network engagement demonstrates that microscale synaptic properties shape macroscale network recruitment. This finding aligns with models highlighting local E/I balance as critical for metastable cortical dynamics (Abeysuriya et al., 2018; Pascoa Dos Santos & Verschure, 2025) and raises the possibility that task-dependent changes in network recruitment, as observed during working memory (Rossi et al., 2023; Rossi et al., 2024) and motor processing (Gohil et al., 2024; Quinn et al., 2018), may reflect underlying adjustments in local timescales or E/I balance (Gao et al., 2020; Manea et al., 2024; Wilson et al., 2022). Future work combining large-scale network analyses with task-based timescale manipulation or targeted manipulation of the regional E/I balance via brain stimulation will be needed to determine whether the effects observed here are specific to the frontal DMN and DAN, or reflect a more general mechanism of network regulation.

Taken together, our findings demonstrate that microscale GABAergic modulation shapes timescales of regional neuronal populations and large-scale network dynamics, highlighting that microscale synaptic properties can drive macroscale cortical organization.

## Methods

This study is part of a larger project that also involves task-based MEG data. The preregistration for the entire experimental protocol can be found at https://doi.org/10.17605/OSF.IO/CMYH2).

### Participants

Sixty male participants took part in the study. Due to excessive noise in the MEG signal, three participants (two of whom had non-magnetic retainers) had to be excluded from analysis, leading to a final sample of 57 participants (Age: 23.58 ± 3.36 years, body weight: 74.9 ± 8.36 kg, body mass index: 22.9 ± 2.12, mean ± SD) with normal or corrected to normal vision (*N* = 13). Most were right-handed (*N* = 52). Due to the pharmacological manipulation, we only included participants that did not have certain preexisting medical conditions, such as psychiatric or neurological disorders (full list of exclusion criteria in Supplement 10). Participants were compensated at a fixed rate plus an additional bonus based on their performance in the experimental tasks (see procedure). All participants were naïve to the purpose of the study and gave written informed consent. All procedures were approved by the local Ethics Committee of the Medical Faculty of the Heinrich Heine University Düsseldorf (reference 2018-211_1). The study was performed in compliance with the Code of Ethics of the World Medical Association (Declaration of Helsinki, 1975).

### Procedure

The study consisted of a medical screening, three pharmacological MEG sessions, and a structural MRI scan. In the screening, it was ensured that participants met the inclusion criteria (full list in Supplement 12). They filled out the Beck’s Depression Inventory (BDI; Beck et al., 1996) and the State-Trait Anxiety Inventory (STAI; Spielberger et al., 1983). Moreover, participants familiarized themselves with the experimental tasks: a value-based decision-making task (VBM), a random-dot motion task (RDM), and a value-based learning task (VBL; experimental tasks will not be explained here but details can be found at https://doi.org/10.17605/OSF.IO/CMYH2). They needed to meet pre-defined performance criteria to participate in the study (see performance-based exclusion criteria in Supplement 10).

The three pharmacological MEG sessions followed identical procedures, only the drug (or placebo) administered differed. In a within-subject, double-blind, cross-over design, participants were given a single oral dose of lorazepam (Tavor®, Pfizer, 1.0mg), D-cycloserine (Thai Meiji Pharmaceutical Co., Ltd., 250 mg), or placebo. MEG sessions were restricted to the morning and separated by at least six days to allow complete washout, based on the drugs’ half-lives (Greenblatt et al., 1979; Patel et al., 2011; Riss et al., 2008; van Berckel et al., 1998). Participants were given a standardized breakfast. Heart rate and blood pressure were measured at three time points per session: before the drug administration, before the MEG measurement, and after the MEG measurement. At the same three time points, participants also filled out the Bond and Lader Visual Analogue Scales to assess drug-related changes in mood (BL-VAS; Bond & Lader, 1974) along with the STAI. Additionally, to control for pharmacological effects on visuomotor processes, participants completed a modified trail-making test before the MEG measurement (Rodewald et al., 2012). The MEG measurement began 2.5 h after drug intake in accordance with the average time for peak plasma concentrations (Greenblatt et al., 1979; Patel et al., 2011; Riss et al., 2008; van Berckel et al., 1998). It started with a 5-minute eyes-open resting state measurement followed by the three experimental tasks (VBM: 30 min, RDM: 50 min, VBL: 20 min). The present analysis focuses on the resting state measurement, and hence the experimental tasks will not be explained here. On the third session, participants were asked to guess the order of drug administration. A total of 26 participants correctly guessed the day they received lorazepam with a mean certainty of *m* = 58.58% (*sd* = 20.44%). Similarly, 25 participants correctly guessed the D-cycloserine session, reporting a certainty of *m* = 56.04% (*sd* = 21.95%). Lastly, a structural MRI scan was recorded on a separate day for each participant.

### Apparatus & resting state measurement

MEG recordings were conducted in a 306-channels whole head MEG system (Elekta Neuromag, Vectorview) with a sampling rate of 1 kHz. Participants’ head position inside the scanner was recorded using four head position indicator (HPI) coils placed at the sides of the forehead and both mastoids. Additionally, eye movement and heart rate were tracked with vertical and horizontal electrooculograms (EOG) and an electrocardiogram (ECG). A full-brain, high-resolution, standard T1-weighted structural magnetic resonance image was acquired from each participant to enable subsequent source reconstruction. To this end, approximately 100 point coordinates were digitized on the participants’ scalp to facilitate accurate co-registration of participants’ MRI scans with MEG data. We recorded 5-minutes eyes-open resting state measurements during which participants were presented with a fixation dot (radius = 0.4 degrees of visual angle; Thaler et al., 2013). It was designed using the PsychoPy software package (version 3.1.5; Peirce, 2007) and presented on a projector (Panasonic PT-D7700E). The screen dimensions were 43 cm x 35 cm with a resolution of 1,280 × 1,024 pixels and a refresh rate of 60 Hz. Participants were placed at a viewing distance of 80 cm.

### Data preprocessing

MEG data was preprocessed with MNE-Python (version 1.0.2; Gramfort et al., 2013) Power line artifacts were removed using the ZapLine method (Rodewald et al., 2012). A 0.1 Hz high-pass filter (FIR filter) was applied. After discarding bad channels based on visual inspection, an independent component analysis (with 60 components) was run (Ablin et al., 2018). Components related to eye movement or heartbeat were removed from the data. Bad time segments were identified through visual inspection and excluded.

### Source reconstruction

Source reconstruction and all subsequent analyses were performed on the gradiometer time series. Forward models were constructed using individual anatomical MRIs with a boundary element model (BEM). Noise covariance matrices were estimated from empty-room recordings acquired at the end of each participant’s MEG session. A linearly constrained minimum variance (LCMV; Sekihara & Nagarajan, 2008) beamformer with a regularization parameter of 5 % was computed using the forward model, noise covariance, and data covariance based on the 5 min measurements. Filters were oriented according to the direction of maximal power. Because MEG is less sensitive to radial sources, the denominator rank was reduced by one at each spatial location.

## Data analyses

### Estimating neuronal timescales

First, we computed the power spectral densities (PSDs) for time series of each vertex using the modified Welch’s method, averaging over time segments with the median instead of the mean to reduce the effect of artefacts (10 s segments with 5 s overlap, Hanning window; Gao et al., 2020; Shafiei et al., 2023). The resulting PSDs were split into periodic and aperiodic activity using the FOOOF toolbox (version 1.0.0; Donoghue et al., 2020). Fitting parameters followed recommendations by Donoghue et al. (2020): A maximum number of 6 peaks were fitted with a peak width between 1 – 6 Hz and a height of at least 0.1 log(Hz). The aperiodic mode was set to “knee” in order to capture the bend in the aperiodic signal. As spectral power settled into a rather steady low level at values higher than 40 Hz, we restricted the frequency range to 0.5 – 40 Hz (Lendner et al., 2024). In cases where no aperiodic knee could be fitted to a power spectrum, it was refitted with the aperiodic mode set to “fixed” (this was the case for 14.21 % out of 1417113 vertices). These fits were disregarded for the analysis of timescales, but for completeness, they were included in the supplementary analyses of drug effects on the aperiodic parameters. All resulting models achieved an average goodness of fit of *R*^2^ = 0.94 (*sd* = 0.03).

The aperiodic activity *L* was estimated using a Lorentzian function characterized by an offset *b*, exponent *x*, and knee *k* (Donoghue et al., 2020).:

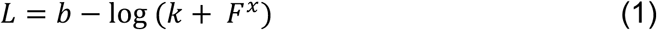

with F denoting frequency (Hz). We then transformed the knee to the knee frequency (*f_k_*) using the exponent (Donoghue et al., 2020):

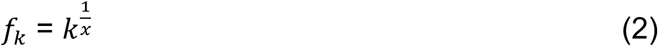

The knee frequency defines the frequency at which the power is no longer constant across frequencies, but instead decreases with increasing frequencies (Donoghue et al., 2020). Finally, using the knee frequency, we could estimate the decay time constant 𝜏 of each vertex (Gao et al., 2020):

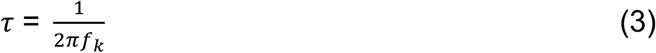

The decay time constant describes the rate at which the autocorrelation function, which quantifies the signal’s self-similarity, decreases over time. For each drug condition, we parcellated the decay time constants into the Desikan-Killiany atlas with 34 cortical regions per hemisphere by computing the median timescale of all vertices in each parcel (Desikan et al., 2006; Shafiei et al., 2023). As a validation check, we correlated our resulting timescale maps with the publicly available intrinsic timescale map (Shafiei et al., 2023; Van Essen et al., 2013).

### Analyzing pharmacological effects on timescales

To investigate whether our timescale map follows the anatomical cortical hierarchy, we correlated our map, for each drug condition, with the T1w/T2w ratio, a structural MRI marker of cortical hierarchy (see statistical analyses; Glasser et al., 2016). Drug effects on neuronal timescales were analyzed per parcel using a permutation test. We further analyzed whether the drug effects on the timescales of each parcel were modulated by their position in the cortical hierarchy. To this end, we implemented a linear mixed-effects model since this approach allows estimating both within-and between-subject variability in accordance with our experimental design (Brown, 2021; Harrison et al., 2018; Jaeger, 2008). Furthermore, we verified that drug effects on neuronal timescales were independent of their effects on control measures (alertness, calmness, contentedness, anxiety, visuomotor processes, heart rate, and blood pressure). For visualization of lorazepam-induced changes in timescales, we computed the percentage change from the placebo to the lorazepam condition.

### TDE-HMM on MEG data

Dynamic large-scale cortical networks were identified using a Time-Delay Embedded Hidden Markov Model (TDE-HMM; Vidaurre et al., 2018) as implemented in osl-dynamics (Gohil, Huang, et al., 2024). This data-driven approach identifies transient brain activity states, each representing distinct patterns of multi-regional oscillatory power and phase synchrony, thereby linking local activity to large-scale network dynamics.

Before the HMM analysis, vertex time courses were parcellated according to the Desikan–Killiany atlas (Desikan et al., 2006). For each parcel, the first principal component across all vertices, assigned to the respective parcel, was extracted to generate a representative parcel-level time series, which was then resampled to 250 Hz. Symmetric multivariate leakage correction (Colclough et al., 2015) and sign-flipping (Vidaurre et al., 2016) were subsequently applied to the 68 parcel time series to minimize spatial leakage and resolve dipole orientation ambiguity. The leakage correction reduced direct and inherited signal spread, as well as “ghost interactions” (Palva et al., 2018), while sign-flipping aligned parcel time-course polarities across participants.

Next, the obtained standardized time series were time-delay embedded (± 7 samples) and reduced to 120 principal components via PCA to capture auto-and cross-covariance structures across regions, while reducing computational costs and avoiding overfitting. On these time series, an HMM was run, inferring 8 states that represent unique covariance patterns reflecting specific spectral power and phase-locking profiles.

Since the HMM inference is randomly initialized, repeated runs with identical parameters can produce slightly different results. Accordingly, the fitting procedure was repeated 10 times, and the 8-state TDE-HMM with the lowest variational free energy is presented in the main analyses. To further ensure robustness across the hyperparameter number of states, all analyses were also replicated using models with 10 and 12 states.

### State-specific spectral descriptions

Participant-specific spectral state descriptions were estimated post hoc by applying a multi-taper approach to the source-reconstructed data, weighted by the TDE-HMM a-posteriori state probabilities (Vidaurre et al., 2016). Time-resolved power and coherence were computed using seven Slepian tapers with a 2 s window, providing ∼2 Hz spectral smoothing. Spectral estimates reflect all occurrences of a given network across visits rather than individual episodes. For visualization, coherence values were thresholded at the 98th percentile to highlight the strongest connections.

### State-specific timescales

State-specific timescales were estimated analogously to the state-specific spectral descriptions but using vertex-level rather than parcel-level time series. To reduce the computational load, Welch’s method was applied to the weighted, source-reconstructed data with the same parameters as in the time-averaged spectral analysis (10 s segments, 5 s overlap, Hanning window). As in the time-averaged analysis, state-specific timescales were derived from vertex-level power spectra, and parcel-level estimates were obtained as the median across all vertices within each parcel. For the visualization of state-specific changes in timescales, we computed the percentage change from the placebo to the lorazepam condition.

### Transition probability analysis

To assess whether transitions between large-scale cortical networks were modulated by the drug condition, transition probability matrices were estimated post hoc from each participant’s hard-classified state time course under placebo and lorazepam. Hard-classified state time courses were obtained by assigning each time point to the most likely HMM state based on the posterior probabilities. For visualization, changes in transition probabilities were expressed as percentage differences between conditions.

## Statistical analyses

### Within-subject comparisons

To assess timescale alterations between pharmacological conditions and between state-specific and time-averaged spectra within participants, two-tailed permutation tests with 10,000 permutations were conducted (Nichols & Holmes, 2002). Resulting *p*-values were corrected for multiple comparisons across parcels, or across parcels and states, using the false discovery rate (FDR) procedure (Genovese et al., 2002). Comparisons were considered significant when FDR-corrected values fell below α = 0.05, both for these analyses and for all other statistical tests.

Drug-related alterations in transition probabilities and state metrics were also assessed using within-participant, two-tailed permutation tests with 10,000 permutations (Nichols & Holmes, 2002). To account for potential dependencies between tests (including the negative correlations arising from row-sum constraints in transition probabilities and the compositional nature of state metrics) statistical significance was determined using maximum t-statistic pooling, as implemented in GLMTOOLs (Quinn et al., 2024), across states, or across states and state transitions, to control for multiple comparisons.

We used a linear mixed-effects model to investigate if the drug effects on the timescales of each parcel were modulated by their position in the cortical hierarchy. The model was implemented in R using the lme4 package (R version 4.2.2; lme4 version 1.1.34; D. Bates et al., 2015; R Development Core Team, 2022). We applied a logarithmic transformation to the outcome variable “timescales” since the distribution was positively skewed. Fixed effects included the main effects and interactions of cortical hierarchy (defined by the T1w/T2w) and drug (lorazepam vs. placebo and D-cycloserine vs. placebo). Following a PCA, the random effect structure includes the main effects of both cortical hierarchy and drug (Douglas Bates et al., 2015). Additionally, as a control analysis, we verified that our drug effects were independent of session order by adding it to the model as a fixed effect. The same approach was implemented for other control measures (mood, anxiety, visuomotor processes, heart rate, and blood pressure) that were significantly modulated by the drug condition.

## Spatial comparisons

To compare our estimated timescales with a publicly available intrinsic timescale brain map (Shafiei et al., 2023; Van Essen et al., 2013) and an MRI-based T1w/T2w ratio brain map (Glasser et al., 2016), we ran Spearman correlations and computed significance using spatial autocorrelation-preserving permutation tests (Alexander-Bloch et al., 2018; Hansen et al., 2022; Markello et al., 2022; Vása & Misic, 2022). Before spatial comparisons, both maps were parcellated into the cortical Desikan-Killiany parcellation (Cammoun et al., 2012; Desikan et al., 2006). Additionally, the publicly available intrinsic timescale map was transformed from 4k to 32k density (Robinson et al., 2018; Robinson et al., 2014).

## Data and code availability

All analysis scripts are publicly available on GitHub: https://github.com/anadiasmaile/GABA-NMDA-timescales. Due to the conditions of our ethics approval, the raw MEG data supporting this study cannot be publicly archived. Instead, we provide the processed derivatives required to reproduce the main analyses reported in the manuscript: https://doi.org/10.17605/OSF.IO/7P4KC.

## Author contributions

- Study conception and design (EO, MIF, HK, GJ), data acquisition (EO, MIF, HK, AADM, ES), data analysis (AADM, OK, EO), interpretation of data for the work (all authors)
- Drafting the article (AADM, OK, EF, GJ), revising it for intellectual content (all authors)
- Approving the final version of the manuscript (all authors)

## Acknowledgements

We thank Hanin Alejel, Helena El Kholy, Judith Geusen, Paul Höchter, Marlene Hüsken, Christina Kalinicenko, Joshua Saal, Kouta Sasaki, Georg Schäfer, and Helena Schmidt for their support during data acquisition. We thank the MRI Core Facility of the University Hospital Düsseldorf, particularly Erika Rädisch and Eric Bechler, for help with the MRI recordings. We further thank Bradley Voytek for his advice regarding aperiodic data analysis and cortical parcellation, and Thomas Pirenne for valuable discussions on data analysis, interpretation, and visualization. Computational infrastructure and support were provided by the Centre for Information and Media Technology at Heinrich Heine University Düsseldorf.

