## Supplementary Material for "Role of GABA and NMDA receptors in shaping cortical timescales and large-scale network dynamics"

| | | |
|---|---|---|
| **Supplement 1:** | **Lorazepam effect on neuronal timescales – significant parcels** | **2** |
| **Supplement 2:** | **Drug effects on aperiodic parameters** | **3** |
| **Supplement 3:** | **Linear mixed effect results of lorazepam effect on timescales** | **5** |
| **Supplement 4:** | **Control measures** | **6** |
| **Supplement 5:** | **Control analyses regarding lorazepam effects on timescales** | **11** |
| **Supplement 6:** | **Repeated measures correlation between lorazepam effects in state-specific timescales and lorazepam effects on time-averaged timescales** | **13** |
| **Supplement 7:** | **Robustness of network-specific timescale findings across different numbers of states** | **14** |
| **Supplement 8:** | **Robustness of lorazepam effects on cortical network dynamics findings across different numbers of states** | **18** |
| **Supplement 9:** | **Large-scale cortical network descriptions are robust across different preprocessing choices** | **20** |
| **Supplement 10:** | **Exclusion criteria** | **23** |
| **Supplement References** | | **25** |

**Supplement 1: Lorazepam effect on neuronal timescales – significant parcels**

**Figure S1: Lorazepam effect on neuronal timescales – significant parcels.** t-statistic derived from a permutation test of the effect of lorazepam on neuronal timescales compared to placebo projected onto the cortical surface. Non-significant parcels are depicted in white. Multiple comparisons across parcels were FDR-corrected. Lorazepam significantly increased neuronal timescales across almost the entire cortex.

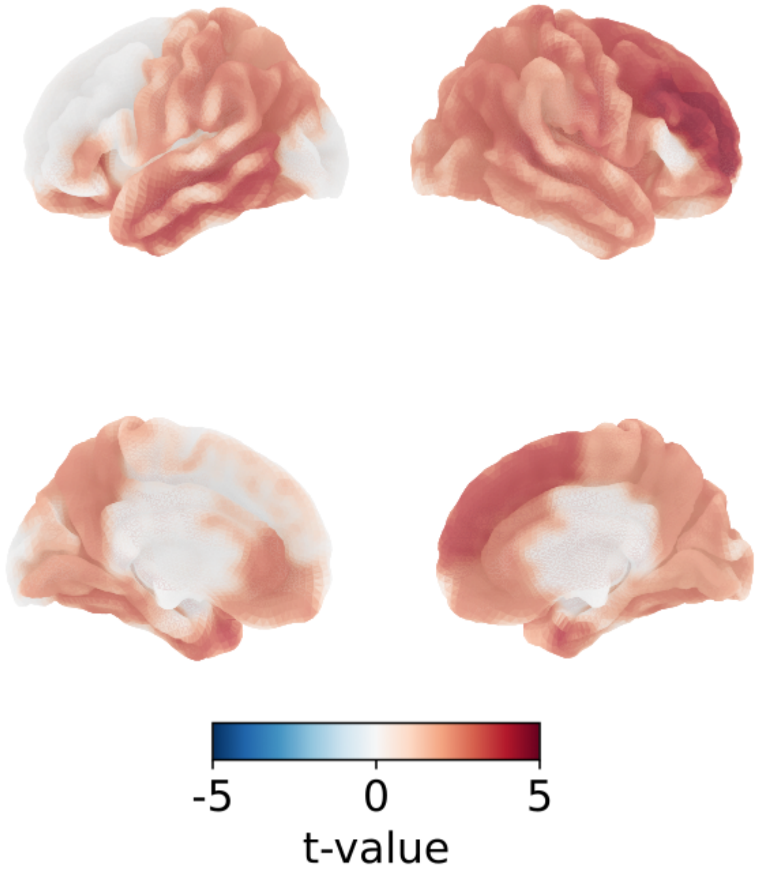

### Supplement 2: Drug effects on aperiodic parameters

In order to assess the effects of lorazepam and D-cycloserine on the aperiodic signal of each cortical region, we performed a two-tailed paired-sample permutation t-test comparing the aperiodic parameters under placebo to both drugs. For each permutation test, we ran 10,000 permutations and applied FDR correction for multiple comparisons.

**Figure S2: Drug effects on aperiodic exponent.** (**a**) Lorazepam significantly decreased the aperiodic exponent in the left entorhinal cortex ($t = -3.33$, $p = 0.034$), left pericalcarine cortex ($t = -3.37$, $p = 0.034$) and the right lateral occipital cortex ($t = -3.66$, $p = 0.014$). Therefore, the slope of the aperiodic signal was significantly flatter in these cortical regions under Lorazepam. (**b**) In contrast, D-cycloserine had no significant effect on the aperiodic exponent.

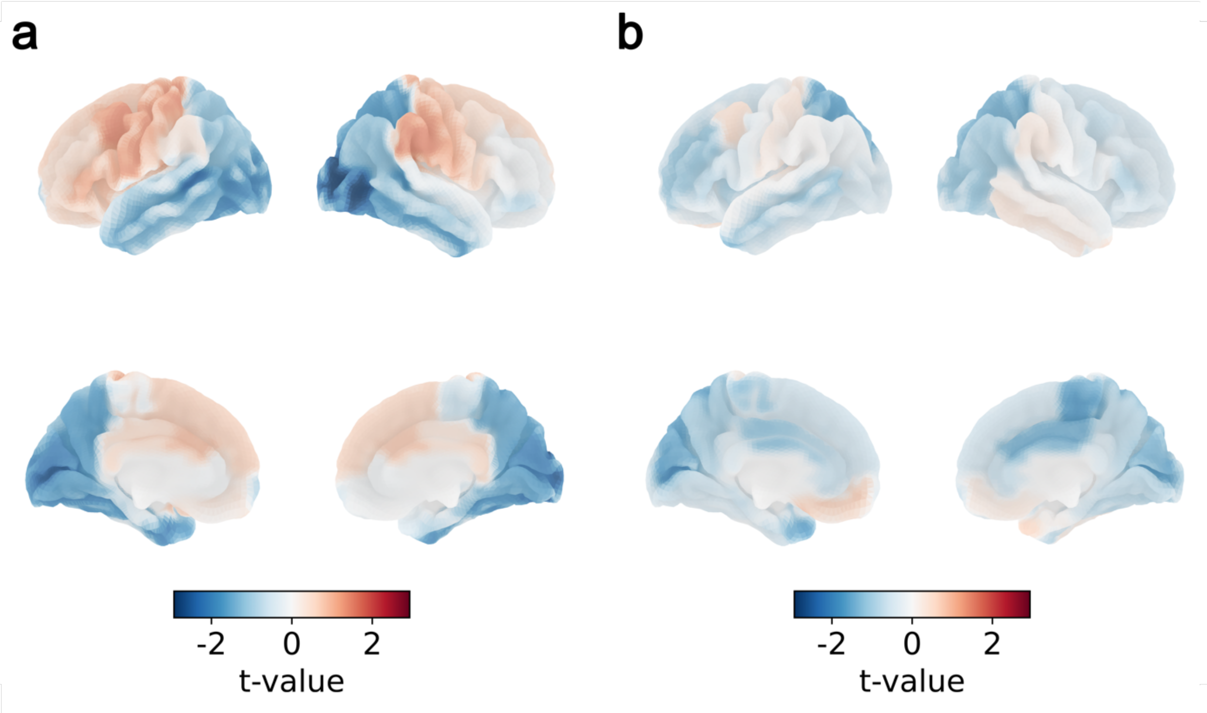

**Figure S3: Drug effects on aperiodic knee frequency.** (**a**) Lorazepam and (**b**) D-cycloserine had no significant effect on the cortical aperiodic knee frequency.

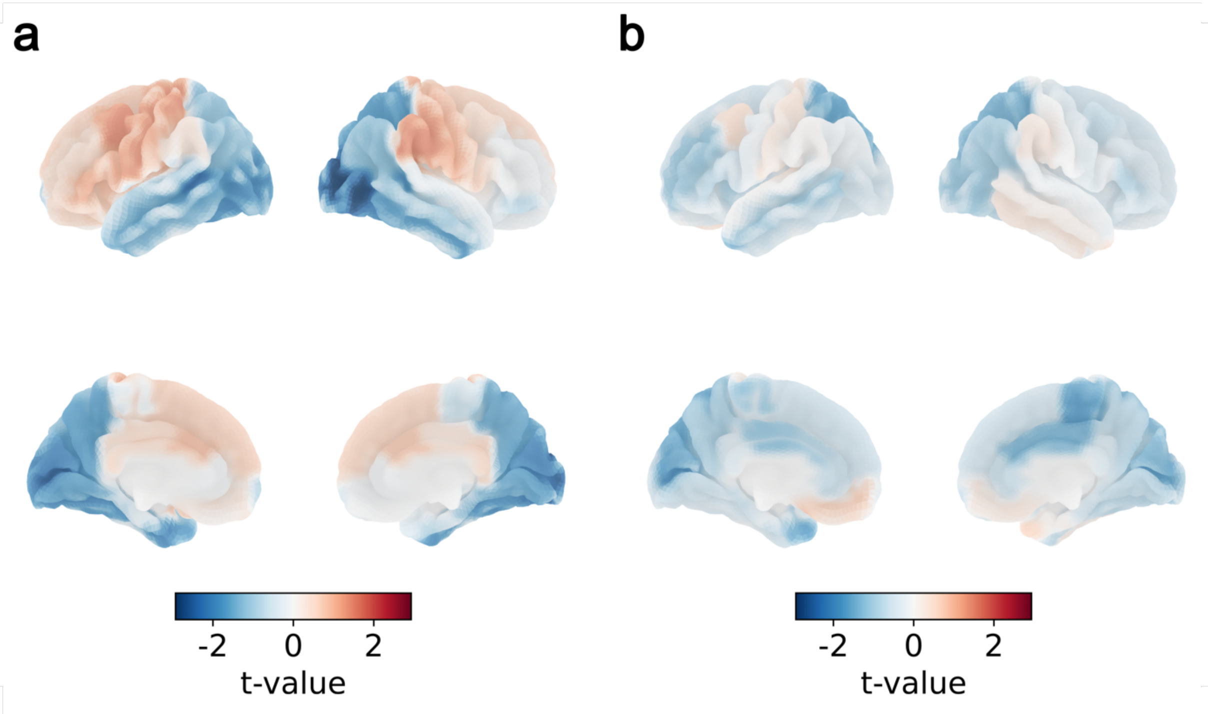


**Figure S4: Drug effects on aperiodic offset.** (**a**) Lorazepam significantly decreased the aperiodic offset in the right lateral occipital cortex ($t$ = -3.33; $p$ = 0.034). (**b**) D-cycloserine had no significant effect on the aperiodic offset.

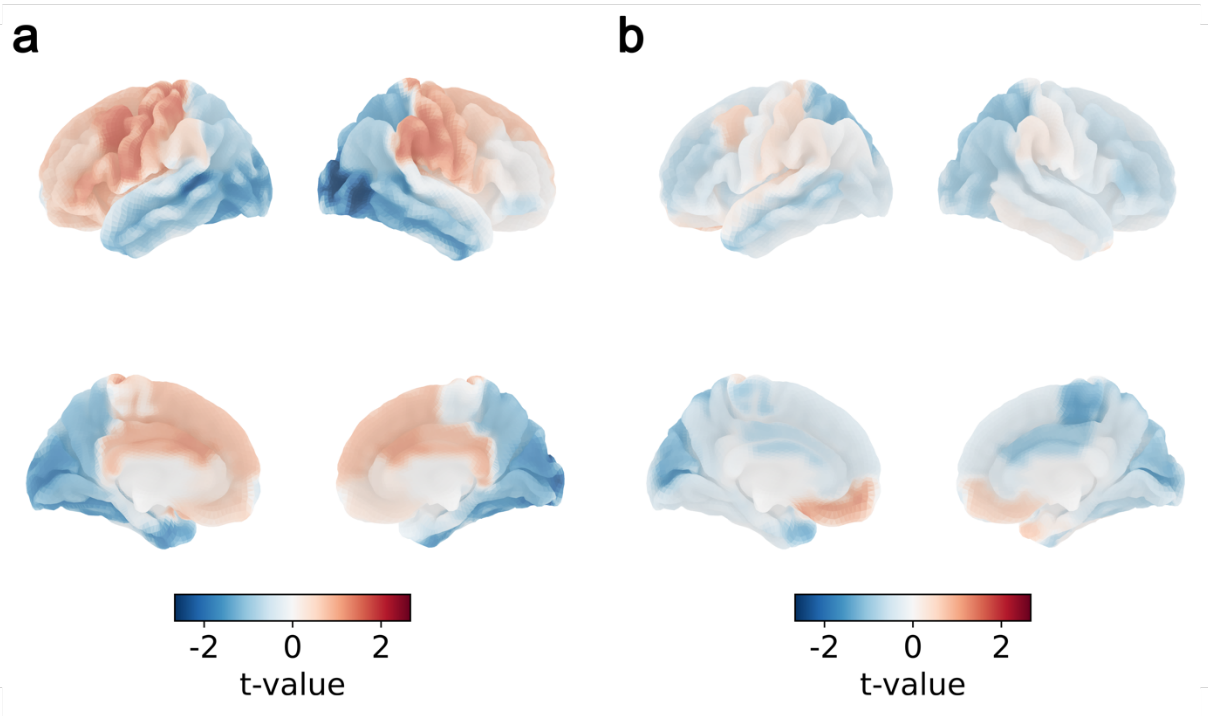

**Supplement 3: Linear mixed effect results of lorazepam effect on timescales**

Full results of the linear mixed model analyzing the joint effect of cortical hierarchy and the drugs on cortical timescales. Timescales significantly increase with decreasing myelination operationalized through the T1w/T2w ratio as a proxy for cortical hierarchy. While D-cycloserine has no significant effect, lorazepam significantly expands the cortical timescales. Additionally, there is a trend effect of the interaction of Lorazepam with the T1wT2w ratio. Thus, the increased timescales through Lorazepam are especially pronounced in frontal regions with typically lower degrees of myelinization.

**Table S1: Linear mixed effect results.** Table includes the β-weight, standard error (SE), *t*-value, degrees of freedom (df) and *p*-value for each fixed effect. Stars represent significant effects based on $p < 0.05$.

| | ***β*-weight** | ***SE*** | **df** | ***t*-value** | ***p*-value** |
|---|---|---|---|---|---|
| Intercept | 0.70 | 0.27 | 69.25 | 2.55 | 0.013 * |
| T1w/T2w | -3.06 | 0.17 | 75.88 | -17.64 | < 0.001 * |
| D-cycloserine | -0.08 | 0.15 | 9952.00 | -0.54 | 0.586 |
| Lorazepam | 0.39 | 0.15 | 5146.00 | 2.55 | 0.011 * |
| T1w/T2w x D-cycloserine | 0.08 | 0.11 | 11180.00 | 0.71 | 0.477 |
| T1w/T2w x Lorazepam | -0.19 | 0.11 | 11190.00 | -1.75 | 0.081 |

### Supplement 4: Control measures

#### Bond and Lader Visual Analogue Scales and State-Trait Anxiety Inventory

Participants' mood and state anxiety was protocolled three times per session: before drug intake (T1), before (T2), and after (T3) the MEG measurement. The resting state measurement was temporally closest to T2. The effects of Lorazepam and D-cycloserine on participants' mood was assessed using linear mixed effects models. Tables include the β-weight, standard error (SE), *t*-value, degrees of freedom (df) and *p*-value for each fixed effect. Stars represent significant effects based on $p < 0.05$.

**Table S2: Alertness score.** Lorazepam significantly reduced alertness at T3. D-cycloserine had no significant effect.

| | ***β*-weight** | ***SE*** | **df** | ***t*-value** | ***p*-value** |
|---|---|---|---|---|---|
| Intercept | 7.65 | 0.23 | 99.88 | 33.00 | < 0.001 * |
| D-cycloserine | 0.15 | 0.18 | 447.05 | 0.87 | 0.384 |
| Lorazepam | 0.01 | 0.18 | 447.00 | 0.08 | 0.939 |
| T2 | -0.30 | 0.18 | 447.00 | -1.68 | 0.093 |
| T3 | -1.05 | 0.18 | 447.00 | -5.96 | < 0.001 * |
| D-cycloserine x T2 | -0.01 | 0.25 | 447.03 | -0.06 | 0.953 |
| Lorazepam x T2 | -0.10 | 0.25 | 447.00 | -0.41 | 0.685 |
| D-cycloserine x T3 | 0.15 | 0.25 | 447.03 | 0.60 | 0.550 |
| Lorazepam x T3 | -0.71 | 0.25 | 447.00 | -2.85 | 0.005 * |

**Table S3: Calmness score.** Lorazepam and D-cycloserine had no significant effect on participants' calmness.

| | ***β*-weight** | ***SE*** | **df** | ***t*-value** | ***p*-value** |
|---|---|---|---|---|---|
| Intercept | 7.96 | 0.19 | 121.46 | 41.12 | < 0.001 * |
| D-cycloserine | -0.06 | 0.17 | 448.00 | -0.36 | 0.723 |
| Lorazepam | 0.18 | 0.17 | 448.00 | 1.05 | 0.293 |
| T2 | 0.39 | 0.17 | 448.00 | 2.31 | 0.021 * |
| T3 | 0.24 | 0.17 | 448.00 | 1.44 | 0.151 |
| D-cycloserine x T2 | -0.04 | 0.24 | 448.00 | -0.15 | 0.879 |
| Lorazepam x T2 | -0.25 | 0.24 | 448.00 | -1.04 | 0.299 |
| D-cycloserine x T3 | 0.01 | 0.24 | 448.00 | 0.04 | 0.972 |
| Lorazepam x T3 | -0.09 | 0.24 | 448.00 | -0.37 | 0.709 |

**Table S4: Contentedness score.** Lorazepam and D-cycloserine had no significant effect on participants' contentedness.

| | *β-weight* | *SE* | df | *t*-value | *p*-value |
|---|---|---|---|---|---|
| Intercept | 8.25 | 0.17 | 92.76 | 48.86 | < 0.001 * |
| D-cycloserine | 0.01 | 0.12 | 446.04 | 0.10 | 0.920 |
| Lorazepam | -0.01 | 0.12 | 446.04 | -0.05 | 0.962 |
| T2 | 0.01 | 0.12 | 446.00 | 0.06 | 0.949 |
| T3 | -0.09 | 0.12 | 446.00 | -0.77 | 0.443 |
| D-cycloserine x T2 | 0.03 | 0.17 | 446.02 | 0.16 | 0.876 |
| Lorazepam x T2 | 0.14 | 0.17 | 446.02 | 0.83 | 0.407 |
| D-cycloserine x T3 | -0.07 | 0.17 | 446.02 | -0.39 | 0.701 |
| Lorazepam x T3 | -0.05 | 0.17 | 446.02 | -0.27 | 0.789 |

**Table S5: State anxiety.** Lorazepam and D-cycloserine had no significant effect on participants' state anxiety.

| | *β-weight* | *SE* | df | *t*-value | *p*-value |
|---|---|---|---|---|---|
| Intercept | 29.47 | 0.74 | 86.20 | 39.95 | < 0.001 * |
| D-cycloserine | 0.57 | 0.49 | 445.05 | 1.16 | 0.245 |
| Lorazepam | 0.39 | 0.49 | 445.05 | 0.80 | 0.422 |
| T2 | 0.06 | 0.49 | 445.09 | 0.12 | 0.905 |
| T3 | 0.76 | 0.49 | 445.09 | 1.54 | 0.124 |
| D-cycloserine x T2 | -0.08 | 0.69 | 445.05 | -0.11 | 0.912 |
| Lorazepam x T2 | -0.04 | 0.69 | 445.05 | -0.06 | 0.953 |
| D-cycloserine x T3 | -0.16 | 0.69 | 445.05 | -0.23 | 0.816 |
| Lorazepam x T3 | 0.17 | 0.69 | 445.05 | 0.25 | 0.802 |

**Blood pressure and heart rate**

Participants' blood pressure and heart rate was measured three times per session: before drug intake (T1), before (T2), and after (T3) the MEG measurement. The resting state measurement was temporally closest to T2. The effects of Lorazepam and D-cycloserine on participants' blood pressure and heart rate was assessed using linear mixed effects models. Tables include the β-weight, standard error (SE), *t*-value, degrees of freedom (df) and *p*-value for each fixed effect. Stars represent significant effects based on $p < 0.05$.

**Table S6: Systolic blood pressure.** Lorazepam significantly reduced the systolic blood pressure at T2. On sessions where D-cycloserine was administered systolic blood pressure was increased across all measuring timepoints. Thus, on these sessions systolic blood pressure was increased independently of the drug administration.

| | ***β*-weight** | ***SE*** | **df** | ***t*-value** | ***p*-value** |
|---|---|---|---|---|---|
| Intercept | 118.79 | 1.49 | 144.02 | 79.83 | < 0.001 * |
| D-cycloserine | 3.89 | 1.40 | 446.01 | 2.79 | 0.006 * |
| Lorazepam | 1.79 | 1.40 | 446.01 | 1.28 | 0.201 |
| T2 | 0.02 | 1.40 | 446.01 | 0.01 | 0.990 |
| T3 | 0.56 | 1.41 | 446.18 | 0.39 | 0.694 |
| D-cycloserine x T2 | -1.88 | 1.98 | 446.01 | -0.95 | 0.343 |
| Lorazepam x T2 | -3.91 | 1.98 | 446.01 | -1.98 | 0.048 * |
| D-cycloserine x T3 | 1.44 | 1.99 | 446.10 | 0.73 | 0.468 |
| Lorazepam x T3 | -0.61 | 1.99 | 446.10 | -0.31 | 0.760 |

**Table S7: Diastolic blood pressure.** Lorazepam had no significant effect on the diastolic blood pressure. On sessions where D-cycloserine was administered diastolic blood pressure was increased across all measuring timepoints. Thus, on these sessions diastolic blood pressure was increased independently of the drug administration.

| | ***β*-weight** | ***SE*** | **df** | ***t*-value** | ***p*-value** |
|---|---|---|---|---|---|
| Intercept | 74.18 | 1.07 | 131.80 | 69.28 | < 0.001 * |
| D-cycloserine | 3.35 | 0.97 | 445.95 | 3.47 | < 0.001 * |
| Lorazepam | 0.02 | 0.97 | 445.95 | 0.02 | 0.986 |
| T2 | -2.84 | 0.97 | 445.95 | -2.94 | 0.003 * |
| T3 | 0.69 | 0.98 | 446.10 | 0.71 | 0.481 |
| D-cycloserine x T2 | -1.81 | 1.37 | 445.95 | -1.32 | 0.186 |
| Lorazepam x T2 | 0.53 | 1.37 | 445.95 | 0.39 | 0.700 |
| D-cycloserine x T3 | -0.35 | 1.37 | 446.03 | -0.26 | 0.797 |
| Lorazepam x T3 | 1.12 | 1.37 | 446.03 | 0.82 | 0.415 |

**Table S8: Heart rate.** Lorazepam significantly increased participants' heart rate at T2 and T3. D-cycloserine had no significant effect.

| | ***β*-weight** | ***SE*** | **df** | ***t*-value** | ***p*-value** |
|---|---|---|---|---|---|
| Intercept | 70.81 | 1.40 | 105.62 | 50.62 | < 0.001 * |
| D-cycloserine | 0.77 | 1.11 | 445.99 | 0.70 | 0.486 |
| Lorazepam | -1.18 | 1.11 | 445.99 | -1.06 | 0.289 |
| T2 | -6.00 | 1.11 | 445.99 | -5.42 | < 0.001 * |
| T3 | -11.22 | 1.12 | 446.09 | -10.03 | < 0.001 * |
| D-cycloserine x T2 | -2.42 | 1.57 | 445.99 | -1.55 | 0.123 |
| Lorazepam x T2 | 3.81 | 1.57 | 445.99 | 2.43 | 0.015 * |
| D-cycloserine x T3 | -1.75 | 1.57 | 446.04 | -1.11 | 0.268 |
| Lorazepam x T3 | 4.24 | 1.57 | 446.04 | 2.69 | 0.007 * |

### Trail-making test

To capture pharmacological effects on visuomotor processes, participants completed the trail-making test before the MEG measurement The effects of Lorazepam and D-cycloserine on participants' visuomotor processes were assessed using linear mixed effects models. The Table includes the β-weight, standard error (SE), *t*-value, degrees of freedom (df) and *p*-value for each fixed effect. Stars represent significant effects based on $p < 0.05$.

**Table S9: Trail-making test.** Lorazepam and D-cycloserine had no significant effects on participants' visuomotor processes

| | ***β*-weight** | ***SE*** | **df** | ***t*-value** | ***p*-value** |
|---|---|---|---|---|---|
| Intercept | 22.79 | 0.88 | 94.43 | 26.04 | < 0.001 * |
| D-cycloserine | 0.86 | 0.76 | 112.00 | 1.13 | 0.260 |
| Lorazepam | 0.96 | 0.76 | 112.00 | 1.27 | 0.206 |

**Supplement 5: Control analyses regarding lorazepam effects on timescales**

Lorazepam significantly reduced participants' systolic blood pressure (Table S6) and alertness (Table S2) and significantly increased their heart rate (Table S8). To control whether the lorazepam induced increase in cortical timescales was independent of the additional effect of the drug, we implemented linear mixed models including these control measures. Control measures were assessed at three times during the testing session. Since the resting state measurement was temporally closest to the timepoint 2 (right before the MEG measurement), linear mixed effect models include the difference of timepoint 2 and 1 (before the drug administration). Additionally, we also verified that the effect of lorazepam was independent of the administration order. Tables include the β-weight, standard error (SE), *t*-value, degrees of freedom (df) and *p*-value for each fixed effect. Stars represent significant effects based on $p < 0.05$.

**Table S11: Linear mixed effect results including systolic blood pressure.** Lorazepam increased cortical timescales independent of its effect on systolic blood pressure.

| | ***β-weight*** | ***SE*** | **df** | ***t*-value** | ***p*-value** |
|---|---|---|---|---|---|
| Intercept | -3.31 | 0.10 | 57.21 | -33.96 | < 0.001 * |
| T1w/T2w | -0.41 | 0.02 | 75.89 | -17.64 | < 0.001 * |
| D-cycloserine | 0.02 | 0.03 | 55.82 | 0.61 | 0.542 |
| Lorazepam | 0.15 | 0.04 | 56.55 | 3.40 | 0.001 * |
| Systolic BP | 0.03 | 0.02 | 55.82 | 1.25 | 0.216 |
| T1w/T2w x D-cycloserine | 0.01 | 0.01 | 1118.00 | 0.71 | 0.478 |
| T1w/T2w x Lorazepam | -0.03 | 0.01 | 1119.00 | -1.75 | 0.081 |
| Systolic BP x D-cycloserine | -0.02 | 0.03 | 67.63 | -0.78 | 0.440 |
| Systolic BP x Lorazepam | -0.11 | 0.05 | 70.13 | -2.35 | 0.021 * |

**Table S12: Linear mixed effect results including alertness score.** Lorazepam increased cortical timescales independent of its effect on the alertness score of the Bond and Lader mood scales.

| | ***β-weight*** | ***SE*** | **df** | ***t*-value** | ***p*-value** |
|---|---|---|---|---|---|
| Intercept | -3.31 | 0.10 | 56.99 | -33.92 | < 0.001 * |
| T1w/T2w | -0.41 | 0.02 | 75.80 | -17.62 | < 0.001 * |
| D-cycloserine | 0.02 | 0.03 | 54.59 | 0.86 | 0.392 |
| Lorazepam | 0.14 | 0.04 | 55.55 | 3.06 | 0.003 * |
| Alertness | 0.00 | 0.02 | 56.71 | -0.08 | 0.935 |
| T1w/T2w x D-cycloserine | 0.01 | 0.01 | 11120.00 | 0.75 | 0.453 |
| T1w/T2w x Lorazepam | -0.03 | 0.01 | 11120.00 | -1.75 | 0.080 |
| Alertness x D-cycloserine | 0.00 | 0.03 | 62.69 | -0.07 | 0.942 |

| | | | | | |
|---|---|---|---|---|---|
| Alertness x Lorazepam | 0.01 | 0.04 | 71.39 | -0.28 | 0.781 |

**Table S13: Linear mixed effect results including heart rate.** Lorazepam increased cortical timescales independent of its effect on the heart rate.

| | ***β*-weight** | ***SE*** | **df** | ***t*-value** | ***p*-value** |
|---|---|---|---|---|---|
| Intercept | -3.31 | 0.10 | 56.90 | -33.92 | < 0.001 * |
| T1w/T2w | -0.41 | 0.02 | 75.87 | -17.62 | < 0.001 * |
| D-cycloserine | 0.02 | 0.03 | 56.43 | 0.86 | 0.392 |
| Lorazepam | 0.14 | 0.05 | 56.28 | 3.06 | 0.003 * |
| Alertness | -0.04 | 0.03 | 56.63 | -0.08 | 0.935 |
| T1w/T2w x D-cycloserine | 0.01 | 0.01 | 11180.00 | 0.75 | 0.453 |
| T1w/T2w x Lorazepam | -0.03 | 0.01 | 11190.00 | -1.75 | 0.080 |
| Alertness x D-cycloserine | 0.04 | 0.03 | 61.89 | -0.07 | 0.942 |
| Alertness x Lorazepam | 0.05 | 0.05 | 82.12 | -0.28 | 0.781 |

**Supplement 6: Repeated measures correlation between lorazepam effects in state-specific timescales and lorazepam effects on time-averaged timescales**

**Figure S5: Repeated-measures correlations between lorazepam-related changes in state-specific and time-averaged timescales.** State-specific timescales for all cortical parcels were correlated with their corresponding time-averaged timescales across parcels, accounting for repeated measures within participants. Each bar represents the Spearman correlation coefficient for a given state. Statistical significance was assessed using permutation tests that shuffled timescale values within participants, with Bonferroni correction applied for multiple comparisons across states. All states except State 1 showed significant correlations, with the frontal default mode network (DMN) exhibiting the strongest association.

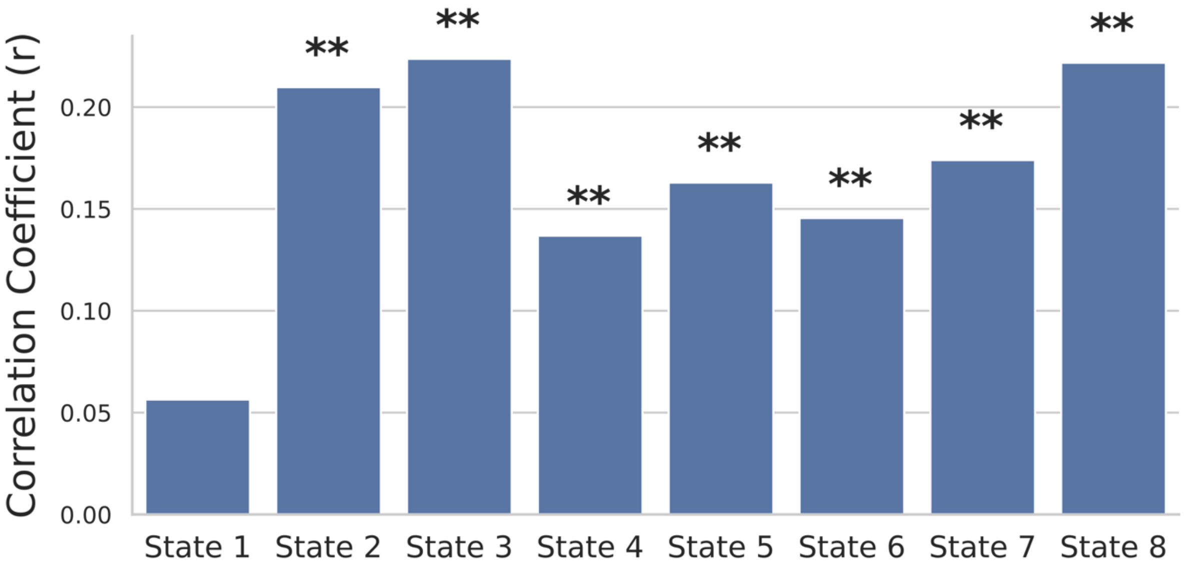

**Supplement 7: Robustness of network-specific timescale findings across different numbers of states**

In the main part of the manuscript, we present findings for a TDE-HMM extracting 8 states. Given that the number of states is a hyperparameter that is a priori specified, we repeated the reported analyses for TDE-HMM fits, inferring 10 and 12 States, to reassure that the main findings of this manuscript are consistent across variations of this hyperparameter. To this end we ran, analogous to the procedure for 8 States, 10 HMMs extracting 10 States and 12 States respectively. For both numbers of states, the fit with the lowest free energy, indicating the best fit to the data, was carried forward for the following analyses. States corresponding to states of interest in the main part of the manuscript (frontal DMN and State 1) were qualitatively identified in the 10- and 12-State fits based on the spectral state description. The matched states are be presented in Supplementary Figure S5.

**Figure S6: Spectral description of networks of interest across TDE-HMM models with different numbers of states.** Spectral descriptions of the frontal default mode network and State 1 obtained from the 8-state TDE-HMM fit reported in the main manuscript (top row) are presented alongside corresponding states from 10-state (middle row) and 12-state (bottom row) fits. For each state, three spectral representations are displayed, averaged across scans from all participants in the placebo condition. Left: Changes in 3-30 Hz power relative to the time-averaged 3-30 Hz power (averaged across all states) projected onto the cortical surface. Top right: State-specific motor cortical power spectrum (black) overlaid with the time-averaged spectrum across all states (grey). Bottom right: Coherence networks in the 2-30 Hz range, thresholded at the 98th percentile to highlight the strongest functional connections. States corresponding to the frontal DMN are outlined in orange.

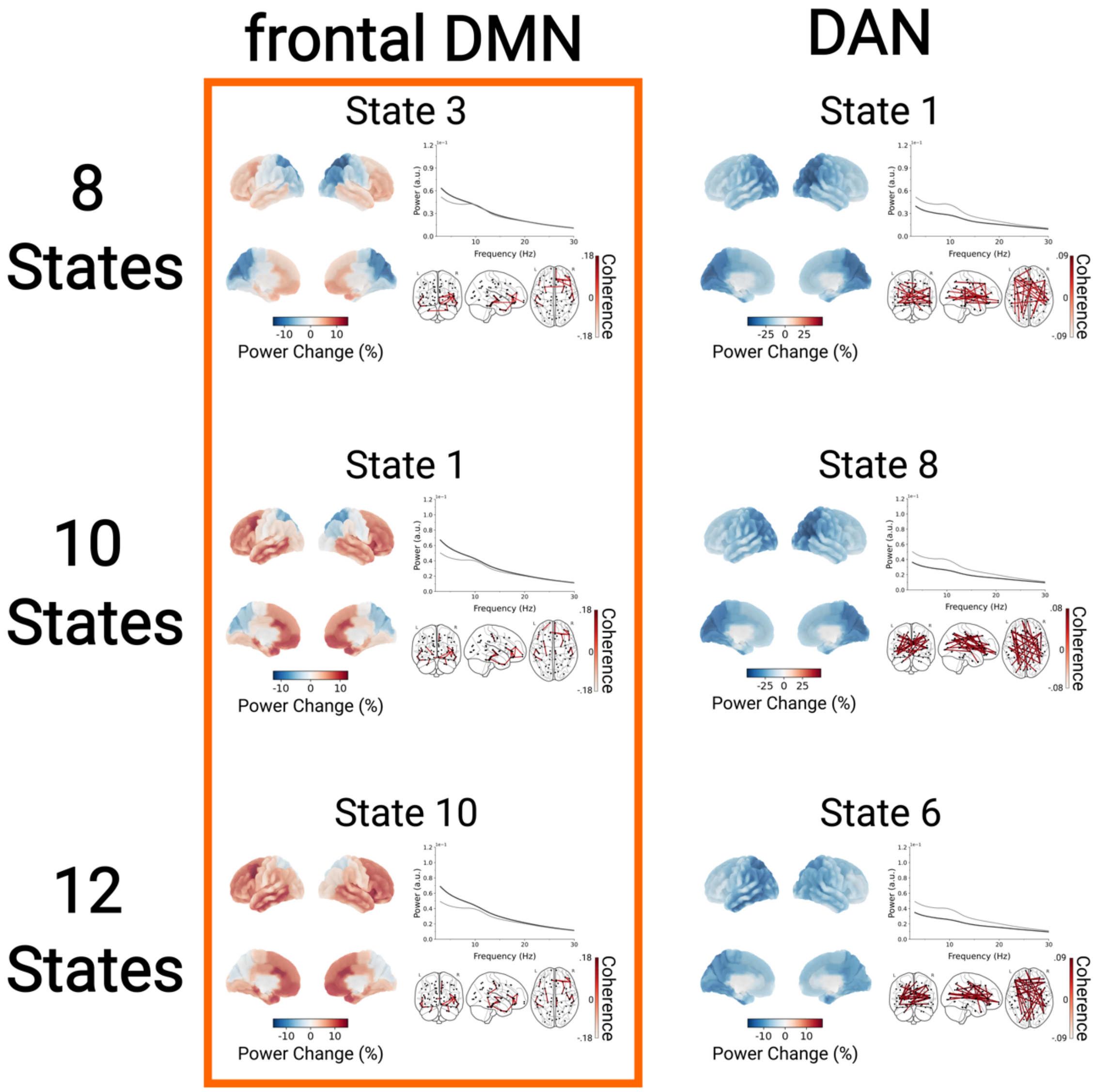


### Variations in timescales across various networks are robust across different numbers of states

Repeating the analyses contrasting network-specific with time-averaged timescales (averaged across all states) produced consistent results for TDE-HMM models with 10 and 12 states (Supplementary Figure S6). In both models, the frontal DMN (10-state HMM: State 1, 52/68 significant parcels, max increase = +6.96% in lingual_lh, $t_{55} = 5.83$, $p < 0.001$; 12-state HMM: State 10, 64/68 parcels, max increase = +14.9% in rostralanteriorcingulate_rh, $t_{55} = 5.21$, $p < 0.001$) and the states corresponding to State 1 from the main analysis (10-state HMM: State 8, 68/68 parcels, max increase = +24.71% in lateraloccipital_lh, $t_{55} = 7.82$, $p < 0.001$; 12-state HMM: State 6, 68/68 parcels, max increase = +22.29% in precuneus_rh, $t_{55} = 7.60$, $p < 0.001$) showed significant increases in timescales during state visits. In contrast, most other states exhibited timescale decreases. As in the main analysis, states corresponding to State 1 displayed strong increases, exceeding those observed in the frontal DMN.

**Figure S7: Network-specific variations in timescales are consistent across models with different numbers of states**. Percentual changes in network-specific timescales relative to the time-averaged scale (averaged across all states) are illustrated for TDE-HMM fits with 10 states (top) and 12 states (bottom). Only cortical parcels showing significant differences between network-specific and time-averaged timescales are displayed ($p < 0.05$, FDR-corrected across parcels and states). The state corresponding to the frontal default mode network in each TDE-HMM fit is outlined in orange.

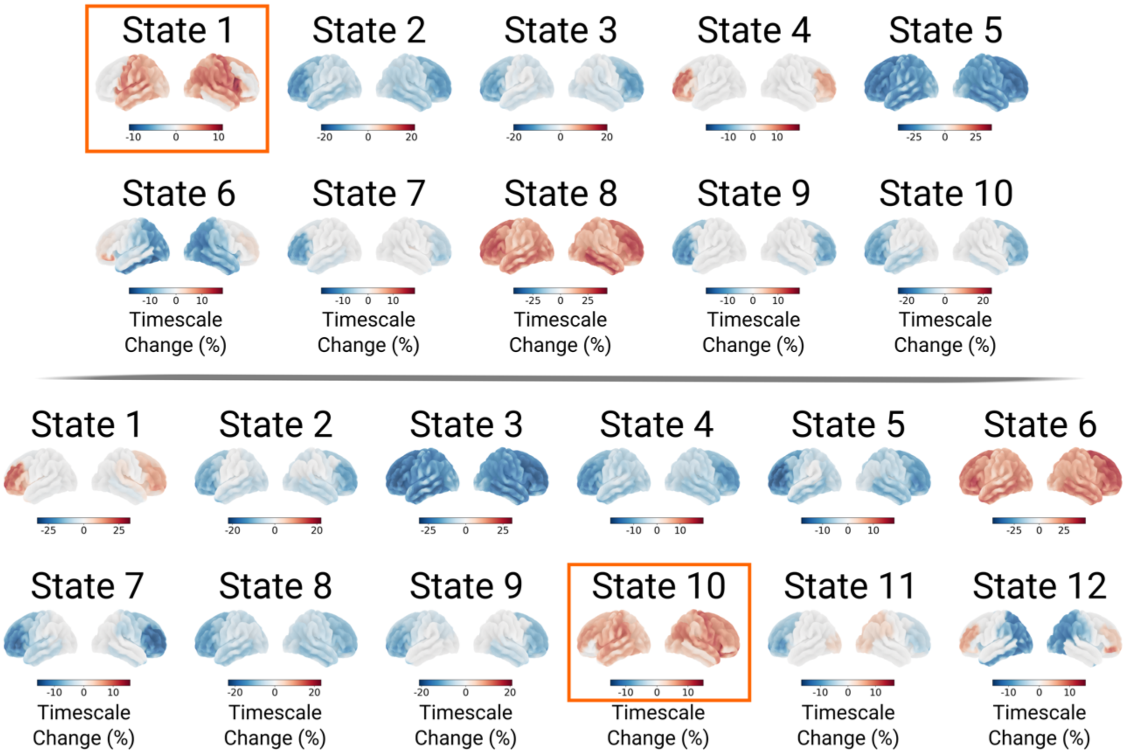


**Lorazepam-related changes in state-specific timescales are robust across different numbers of states**

Repeating the analyses contrasting network-specific timescales between the lorazepam and placebo conditions yielded consistent results across TDE-HMM models with 10 and 12 states (Supplementary Figure S7). Both models revealed distinct patterns of lorazepam-related increases and decreases in timescales across networks. In line with the main findings, lorazepam significantly increased timescales during visits to the frontal DMN (10-state: State 1, 44/68 significant parcels, max increase = +8.96% in lateraloccipital_rh, $t_{55}$ = 4.01, $p$ < 0.001; 12-state: State 10, 36/68 parcels, max increase = +14.46% in fusiform_rh, $t_{55}$ = 4.69, $p$ < 0.001) and in the networks corresponding to State 1 from the main analysis (10-state: State 8, 7/68 parcels, max increase = +9.84% in inferiorparietal_rh, $t_{55}$ = 3.40, $p$ = 0.001; 12-state: State 6, 6/68 parcels, max increase = +13.61% in inferiorparietal_rh, $t_{55}$ = 3.34, $p$ = 0.001). Although less pronounced, the frontal DMN still exhibited greater timescale increases than most other states in both models.

**Figure S8: Lorazepam-related modulations of cortical network-specific timescales are consistent across models with different numbers of states.**
Percentual change in network-specific timescales after lorazepam intake relative to the placebo condition for TDE-HMM fits with 10 states (top) and 12 states (bottom). Only parcels showing significant lorazepam-related differences are displayed on the

cortical surface. Significance was assessed for each state and parcel using within-subject GLMs comparing lorazepam and placebo conditions, with FDR correction applied across all parcels and states ($p < 0.05$). The state corresponding to the frontal default mode network in each TDE-HMM fit is outlined in orange.

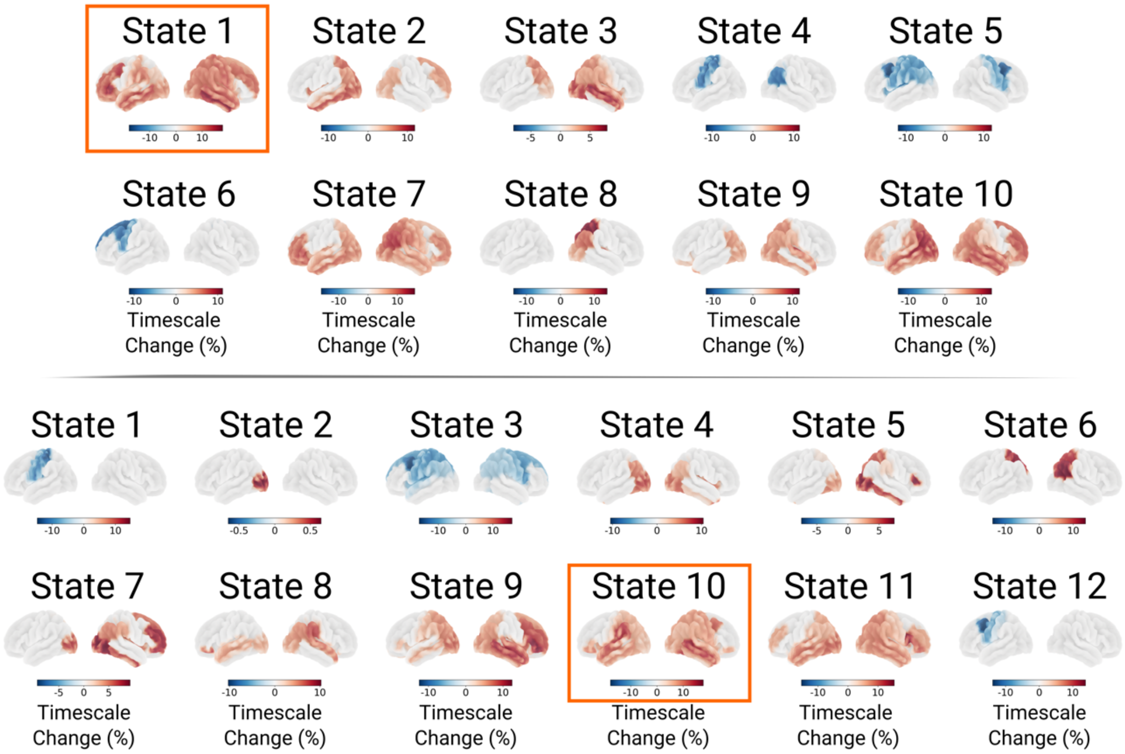

### Supplement 8: Robustness of lorazepam effects on cortical network dynamics findings across different numbers of states

#### Lorazepam-related increases in frontal DMN occurrence probability are robust across different numbers of states

Repeating the analyses of state metrics describing network dynamics produced consistent results across TDE-HMM models with 10 and 12 states (Supplementary Figure S9). Fractional occupancy of the frontal default mode network (DMN) was significantly increased following lorazepam administration in both models (10-state HMM: State 1, $t_{55} = 3.23$, $p = 0.02$; 12-state HMM: State 10, $t_{55} = 3.05$, $p = 0.04$). Conversely, the state corresponding to State 1 from the main analysis revealed a significant reduction in occurrence probability (10-state HMM: State 8, $t_{55} = -3.35$, $p = 0.01$; 12-state HMM: State 6, $t_{55} = -3.28$, $p = 0.02$). Increases in state rates of the frontal DMN (State 1, $t_{55} = 3.11$, $p = .03$) and decreases in lifetimes of the state corresponding to State 1 from the main analysis (State 8, $t_{55} = -2.96$, $p = .03$) were observed only in the 10-state TDE-HMM, but not in the 12-state model. Together, these findings suggest that lorazepam-related increases in frontal DMN occupancy and decreases in State 1 occurrence probability are robust across models with different numbers of inferred states, while the precise mechanisms underlying these changes (e.g., alterations in lifetimes or state rates) appear less consistent.

**Figure S9: Lorazepam-related increases in frontal DMN occupancy are consistent across TDE-HMM models with different numbers of states.** Lorazepam-related changes in state metrics of the frontal default mode network and the state corresponding to State 1 in the main manuscript are shown for TDE-HMM fits with 10 states (**a**) and 12 states (**b**). **(a**) In the 10-state model, State 1 corresponds to the frontal DMN and State 8 to State 1 in the main manuscript. Each dot represents an individual participant's metric in the respective drug condition. Condition contrasts were assessed using maximum t-statistic permutation tests, corrected for multiple comparisons across states. Lorazepam significantly increased fractional occupancy ($t_{55}$ = 3.23, $p$ = 0.02) and state rates ($t_{55}$ =3.11, $p$ = .03) for the frontal DMN, while fractional occupancy ($t_{55}$ = –3.35, $p$ = 0.01) and lifetimes ($t_{55}$ = -2.96, $p$ = .03) for State 8 were reduced. (**b**) In the 12-state model, State 10 corresponds to the frontal DMN and State 6 to State 1 in the main manuscript. Lorazepam increased fractional occupancy for the frontal DMN ($t_{55}$ = 3.05, $p$ = 0.04) and reduced fractional occupancy for State 6 ($t_{55}$ = –3.28, $p$ = 0.02). Asterisks denote significance levels: $p$ < 0.01 (**), $p$ < 0.05 (*).

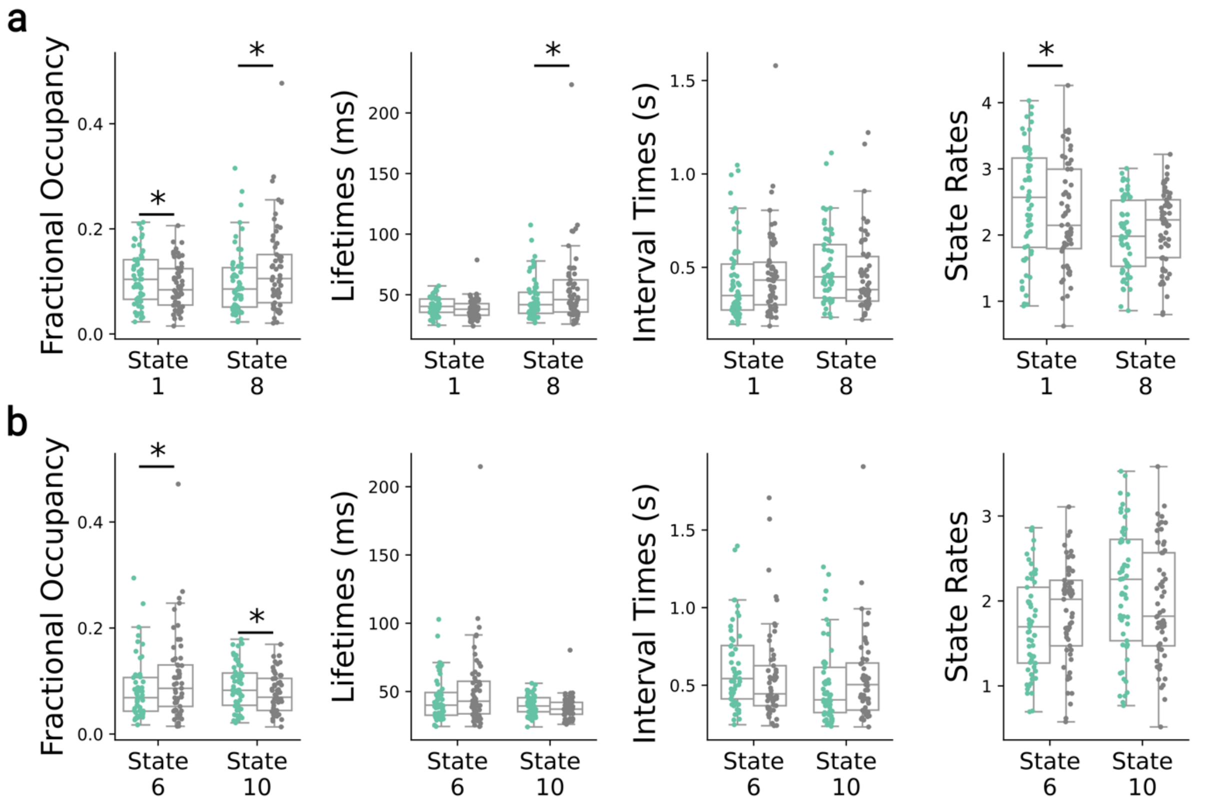

### Supplement 9: Large-scale cortical network descriptions are robust across different preprocessing choices

Given that the preprocessing procedures used in the main analyses deviated in several aspects (e.g., no use vs. application of MaxFilter, use of the OSL LCMV beamformer vs. the MNE beamformer implementation, and parcellation using the Desikan–Killiany atlas vs. the Glasser52 parcellation) from pipelines commonly used in TDE-HMM analyses, we tested whether the network descriptions reported in the manuscript could be replicated using a standard preprocessing pipeline. To this end, the preprocessing was repeated following the pipeline adopted in previous studies running TDE-HMM analyses.[1–4] For a detailed description of the preprocessing pipeline, see Kohl et al. 2025.[1]

#### Overview of the alternative preprocessing pipeline

In short, the raw MEG data were processed using MNE-Python's Maxwell filtering pipeline, including automatic bad-channel detection and temporal signal-space separation (tSSS) for external noise suppression.[5,6] Data were bandpass filtered (0.5–125 Hz), notch-filtered at 50 Hz and 100 Hz, and resampled to 250 Hz. Artefactual segments were excluded using a window-based Generalised ESD test.[7] Cardiac and ocular artefacts were removed via signal-space projection (SSP).[8] Data quality was verified through spectral inspection, and two participants were excluded due to poor data quality.

MEG-MRI co-registration was performed using RHINO, excluding nasal points due to the limited MRI field of view. Source reconstruction employed broadband (1–45 Hz) filtered, sensor-normalised data using an LCMV beamformer (rank = 60),[9,10] and results were projected onto an 8 mm MNI152 template. Source-level signals were parcellated using a reduced Glasser atlas,[11,12] extracting the first principal component per parcel.

Before HMM analysis, symmetrical multivariate leakage correction and sign-flipping were applied to minimise spatial leakage and align dipole polarities across participants.

#### Comparison of large-scale cortical network descriptions

The TDE-HMM fitting procedure was performed 10 times on the alternatively pre-processed data, and the fit with the lowest free energy, indicating the best fit to the data, was carried forward for the comparison of the TDE-HMM fits on the differently pre-processed datasets.

To compare large-scale cortical network descriptions obtained from TDE-HMMs, extracting 8 states from the differently pre-processed datasets, were matched by firstly calculating the correlation distance (1 – Person Correlation) between all states' fractional occupancies in the two data sets. Afterwards, the best assignment of states between the two different HMM fits was inferred with the linear_sum_assignment function in scipy, finding the solution with the lowest overall distance between all assigned pairs.

Large-scale cortical network descriptions of the matched states are presented in Supplementary Figure S4. Qualitative comparisons of the matched states indicate that large-scale cortical network descriptions derived from differently preprocessed datasets exhibit only minor discrepancies. Overall, the patterns are very similar, enabling clear state-to-state matching.

To compare large-scale cortical network descriptions obtained from TDE-HMMs that extracted 8 states from the differently pre-processed datasets, states were matched across HMM fits by first computing the correlation distance (1 - Pearson correlation)

between the fractional occupancies of all states in the two datasets. Subsequently, the optimal one-to-one assignment of states between the two HMM fits was determined using the linear_sum_assignment function from scipy, which identifies the pairing with the lowest overall distance between all matched states.
Large-scale cortical network descriptions of the matched states are presented in Supplementary Figure S4. Qualitative comparison of the matched states revealed that, while minor discrepancies were observed, the large-scale network patterns inferred from the differently preprocessed datasets were overall highly consistent. Importantly, both TDE-HMM solutions identified a network characterized by increased wideband power in frontal and temporal regions, accompanied by power reductions in posterior areas and pronounced low-frequency power increases in the average power spectrum. These observations indicate that this network is reliably represented across both preprocessing variants, suggesting that its identification is robust to differences in the data preparation.

**Figure S10: Comparison of large-scale cortical network descriptions obtained from different preprocessing pipelines.** Spectral representations of cortical networks derived from the preprocessing pipeline described in the main manuscript are depicted on the left, alongside corresponding network representations obtained using preprocessing approaches commonly applied in TDE-HMM studies. States across TDE-HMM fits were matched by minimising the correlation distance between all state pairs (see above). The spectral profiles of matched states showed minor differences but were overall very similar, indicating that network estimates were robust to preprocessing choices. For a detailed description of the spectral state descriptions, refer to Figure 3 in the main text.

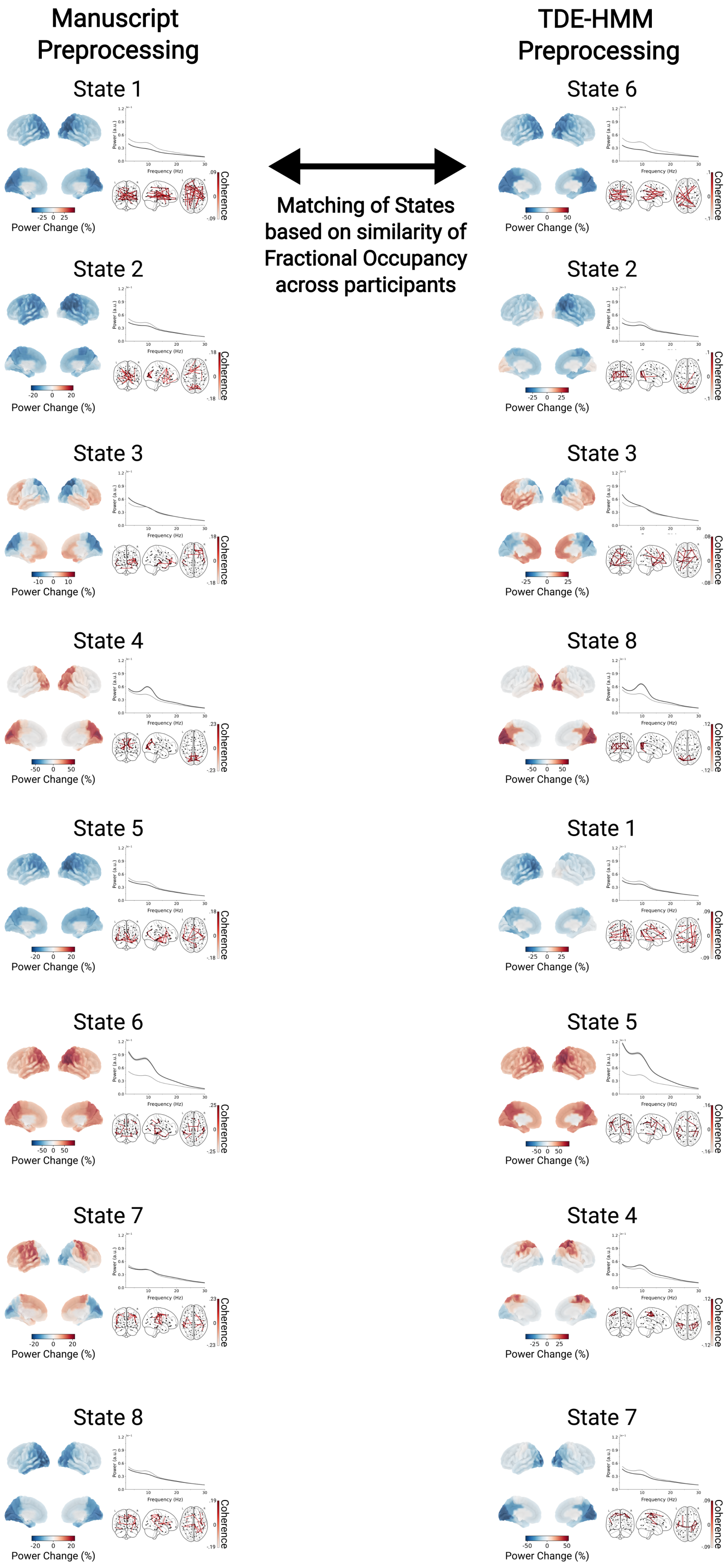
Manuscript Preprocessing
TDE-HMM Preprocessing
Matching of States based on similarity of Fractional Occupancy across participants
State 1
State 6
State 2
State 2
State 3
State 3
State 4
State 8
State 5
State 1
State 6
State 5
State 7
State 4
State 8
State 7
Power Change (%)
Coherence

**Supplement 10: Exclusion criteria**

- Female
- Age < 18 or > 35
- Body weight < 60 or > 90
- BMI < 18 or > 28
- Metal implants (e.g. Retainer) that interfered with the MEG signal or pacemaker
- Claustrophobia
- Smoking > 5 cigarettes per day
- Consumption of > 12 units alcohol per week
- (History of) significant drug abuse (cannabis, amphetamine, cocaine, ecstasy, barbiturate, tranquilizer, opiate, psychodelia, etc.). Significant is defined as:
    1. Use during the last month
    2. exceeding recreational use (> 5 times in lifetime) except or cannabis
    3. frequent use (on average more than once a month) of cannabis
- High blood pressure (systolic > 150 or diastolic > 100)
- Head injury in the past that led to loss of consciousness, or loss of consciousness with an unexplained cause in the past
- (History of) psychiatric or neurological treatment
- Beck's Depression Inventory score > 12
- Medication use interfering with lorazepam or D-cycloserine use
- Epilepsy
- Glaucoma
- Thyroid dysfunction
- Stomach or duodenal ulcer
- Stenosis in gastrointestinal tract
- (History of) gastrointestinal bleedings
- Megacolon
- Ileus
- Prostate adenoma with residual urine formation
- Diabetes
- Asthma
- Impaired liver or kidney function
- Disorders that can lead to tachycardia
- Disorders of cardiovascular system (e.g. arrythmia, high blood pressure, hemophilia)
- Performance-based exclusion:
    - VBM: choice of high expected value option < 55%
    - RDM: accuracy below chance-level, no parametric modulation as a function of coherence level
    - VBL: choice of high-expected value option and high-likelihood option < 55%
